# Insights into the structure and function of the human organic anion transporter 1 in lipid bilayer membranes

**DOI:** 10.1101/2022.01.10.475390

**Authors:** Angelika Janaszkiewicz, Ágota Tóth, Quentin Faucher, Marving Martin, Benjamin Chantemargue, Chantal Barin-Le Guellec, Pierre Marquet, Florent Di Meo

**Affiliations:** INSERM U1248 Pharmacology & Transplantation, Univ. Limoges, CBRS, 2 rue du prof. Descottes, F-87000 Limoges, France; InSiliBio, 1 avenue d’Ester, Ester Technopôle, F-87000 Limoges, France; CHU de Tours, 2 Boulevard Tonnellé, F-37044 Tours, France; Department of Pharmacology and Toxicology, CHU Limoges, F-87000 Limoges, France

**Keywords:** Membrane Transporters, Structural Pharmacology, Molecular Dynamics, Protein-lipid interactions, Major Facilitator Superfamily, Organic Anion Transporter 1

## Abstract

The human SLC22A6/OAT1 plays an important role in the elimination of a broad range of endogenous substances and xenobiotics thus attracting attention from the pharmacological community. Furthermore, OAT1 is also involved in key physiological events such as the remote inter-organ communication. Despite its significance, the knowledge about *h*OAT1 structure and the transport mechanism at the atomic level remains fragmented owing to the lack of resolved structures. By means of protein-threading modeling refined by μs-scaled Molecular Dynamics simulations, the present study provides the first robust model of *h*OAT1 in outward-facing conformation. Taking advantage of the AlphaFold 2 predicted structure of *h*OAT1 in inward-facing conformation, we here provide the essential structural and functional features comparing both states. The intracellular motifs conserved among Major Facilitator Superfamily members create a so-called “charge-relay system” that works as molecular switches modulating the conformation. The principal element of the event points at interactions of charged residues that appear crucial for the transporter dynamics and function. Moreover, *h*OAT1 model was embedded in different lipid bilayer membranes highlighting the crucial structural dependence on lipid-protein interactions. MD simulations supported the pivotal role of phosphatidylethanolamine components to the protein conformation stability. The present model is made available to decipher the impact of any observed polymorphism and mutation on drug transport as well as to understand substrate binding modes.

## 1. Introduction

Major Facilitator Superfamily (MFS) proteins belong to the SoLute Carrier (SLC) superfamily, one of the most important classes of membrane transporters. They can translocate a broad range of endogenous compounds and xenobiotics across cell membranes and play important pharmacological and physiological roles^1,2^. MFS transporters can affect drug pharmacokinetics by modulating absorption, distribution and elimination^3^ since they are involved in cellular influx or efflux. Understanding MFS transporter functions and kinetics is of particular importance to decipher how do they modulate the local pharmacokinetics *i.e*., local drug concentration at the target sites, whether linked with xenobiotic journey and/or therapeutic/adverse effects. This is particularly relevant since, over the past years, growing interest has been paid to local xenobiotic bioavailability (*i.e*., at the intracellular scale)^4^ which can help fulfill the gap between systemic and cellular-scaled pharmacological investigations^5^.

From the physiological point of view, MFS transporters also play an essential role in maintaining homeostasis at the systemic and cellular scales. MFS transporters are involved in cellular nutrient disposition^1,2^ (*e.g*., sugar porters including Glucose transporters – GLUTs – family) as well as in detoxification processes^6,7^ (*e.g*., Organic Anion Transporter family). By modulating body fluid and tissue concentrations of a broad range of specific endo/exogenous molecules, MFS transporters might even drive hormone-independent remote inter-organ communications^8^. The so-called “remote sensing signaling theory”^2^ is key to rationalize the remote modulation of transporter expressions or functions in distant organs as already suggested for SLC^8,9^ and ATP-Binding Cassette (ABC) transporters, in physiology but also in pharmacology^10^.

Several studies have provided evidence in favor of the central role of human Organic Anion Transporter 1 (*SLC22A6*/OAT1) in this dual physiology/pharmacology context^11^. Originally known as the New Kidney Transporter (NKT), *h*OAT1 is a multi-specific transporter mostly expressed in kidneys^12^, at the basolateral membrane of proximal tubular cells (PTC) where it participates in the substrate uptake phase of blood-urine PTC exchanges^13^. *h*OAT1 transports mostly anionic compounds, including xenobiotics such as antiviral acyclic nucleoside phosphonates (*e.g*., tenofovir, adefovir)^14^, endogenous compounds and metabolites (*e.g*., mono- and di-carboxylates) including uremic toxins, especially protein-bound uremic toxins (*e.g*., indoxyl sulfate, *p*-cresol sulfate)^6,7,15,16^. Therefore, *h*OAT1 dysfunctions are not only associated with the impairment of xenobiotic elimination, but also with pathophysiological conditions owing to increased systemic retention of uremic toxin such as in Chronic Kidney Disease^6,7,16^. Furthermore, a large diversity of substrates may compete between them for *h*OAT1 transport^2^. Likewise, several xenobiotics act as *h*OAT1 inhibitors and affect *h*OAT1-mediated detoxification processes^16^. These competition events can also impair drug therapeutic efficacy or lead to adverse effects^15,17^. *h*OAT1 impairment has long been assumed to have a limited impact owing to the redundant expression of *h*OAT3, with which it has a significant substrate overlap. However, this importance of *h*OAT1/*h*OAT3 duality should not be overestimated owing to the recently described substrate selectivity regarding metabolites^18^. This explains the recommendation from the International Transporter Consortium about the evaluation of *h*OAT1 activity in drug discovery^19,20^, followed by the Food and Drug Administration^21^, the European Medicine Agency and the Japan Pharmaceutical and Medical Devices Agency, at least in term of inhibition studies^22,23^.

Despite the great importance of *h*OAT1 in terms of xenobiotic renal clearance, knowledge about the transport mechanism remains fragmented. *h*OAT1 is an antiporter, translocating substrates from blood to the intracellular compartment in exchange for at least one *α*-ketoglutarate (αKG)^13^. Substrate translocation is expected to be driven by αKG concentration which is governed by both the Na^+^/dicarboxylate transporter (*SLC13A3*/NaDC3) and intracellular metabolism^12^. It is worth mentioning that *h*NaDC3 activity in PTCs is strongly related to Na^+^/K^+^-ATPase, leading to a “tertiary” active transport involving the Na^+^/K^+^-ATPase – NaDC3 – OAT1 triad^12^. At the nanoscale, substrate translocation is expected to follow alternating access involving at least three conformational states, namely the outward-facing (OF), occluded and open inward-facing (IF) states^3^. Regarding the unknown folding of *h*OAT1, only two structural models of *h*OAT1 in lipid bilayer membranes have been reported so far. They were obtained by refining homology models with short 100ns+ molecular dynamics (MD) simulations^24,25^. Both models adopted the IF state, using bacterial *E. coli* Glycerol-3-phosphate Transporter (GlpT) resolved structure as the initial template^26^. A high-confidence IF model was recently released using the machine-learning structural prediction tool AlphaFold 2 (AF2)^27^.

Even though key residues can be identified from computational as well as experimental studies the dynamic and atomic features of *h*OAT1 structure still remain unclear^18,24,28,29^ (see Supplementary Table S1 for details). The absence of a robust *h*OAT1 OF model precludes the thorough atomistic rationalization of substrate binding events as well as the investigation of lipid-protein interactions, which have been shown to be of major importance for several MFS transporters and other membrane proteins by either experimental or computational techniques^30–33^. Furthermore, within the frameworks of pharmacogenetics (PGx), atomic-scaled and dynamic pictures of MFS transporters enable the investigation of the structural (and possible functional) modifications arising from Single Nucleotide Polymorphisms (SNPs) or rare pharmacogenetic mutations. In the present study, we propose a protein threading-based model of the missing OF state of *h*OAT1. Microsecond-long MD simulations of *h*OAT1 inserted in several lipid bilayer membranes were performed in order to: (i) refine the initial protein threading static model; (ii) explore the local conformational space of *h*OAT1; and (iii) assess lipid-protein interactions. Topology and structure of the proposed models were carefully analyzed and systematically confronted to AF2 model, as well as to experimental observations. We propose here mechanistic and structural insights into *h*OAT1 transport including the role of lipid-protein interactions.

## 2. Methods

### 2.1. Putative structure of *h*OAT1 in outward-facing state

The amino acid sequence of *h*OAT1 was obtained from UniProt database^34^ with the accession number Q4U2R8, using isoform 1 as the canonical sequence. The initial three-dimensional model of wild-type *h*OAT1 was achieved using the automated protein structure prediction tool I-TASSER webserver^35^. Three relevant resolved MFS proteins were identified as templates, namely *h*GLUT3 (PDB ID: 5C65, 2.65Å resolution)^36^, *r*GLUT5 (PDB ID: 4YBQ, 3.27Å resolution)^37^ and XylE (PDB ID: 4GBY, 2.81Å resolution)^38^, for which sequence identities and similarities are reported in Supplementary Table S2 as well as sequence alignments in Supplementary Figure S1. It is worth mentioning that the initial I-TASSER *h*OAT1 model exhibited a salt bridge between Asp112 and Thr540. This would lead to an implausible direct polar interaction between the extracellular loop (ECL) 1 and the intracellular C-terminal domain in the lipid bilayer. This artifact was thus corrected by means of steered MD simulations in which both Asp112 and Thr540 were pulled apart from each other. A steered MD simulation in pure 1-palmitoyl-2-oleoyl-sn-glycero-3-phosphocholine (POPC) lipid bilayer membrane (see section 2.2. regarding the embedding procedure used) was first performed to smoothly increase the distance between Asp112 and lipid bilayer membrane centers-of-mass (COM) while maintaining Thr540 by positional restraints. Then, the distance between Thr540 and the lipid bilayer membrane COMs was increased. Both simulations were carried out for 2 ns, applying a restraint force constant potential of 35 kcal/mol/Å^2^ in the z-direction and a pulling velocity of 10 Å/ns. In order to improve our initial model, μs-scaled MD simulations were performed including the surrounding environment (*i.e*., lipid bilayer membrane, water and ions) following an approach similar to that previously used for the human multidrug resistance-associated protein 4 (*ABCC4*/MRP4)^39^. Besides, the released AF2 *h*OAT1 IF model was also considered for MD simulations.

### 2.2. Model preparation for MD simulations

Protonation states of charged residues were assigned, using the PROPKA software^40^, at pH = 7.4. Special attention was paid to histidines to which protonation states were assigned in accordance with their calculated pKa as well as potential H-bond networks with surrounding residues by visual inspection. The *ε*-protonated state was used for His47, His130, His217, His246, His249 and His546, the δ-protonated state for His48, His275 and His337, while the cationic double ε/δ-protonated state was assigned to His34. The C-terminal domain (549-563) was cut out of the model to avoid unexpected interactions owing to its high flexibility. The resulting *h*OAT1 model was then embedded in lipid bilayer membranes using the CHARMM-GUI membrane builder tool^41^. Four different POPC-based lipid bilayer membranes were considered, representing different molecular ratios of 1-palmitoyl-2-oleoyl-sn-glycero-3-phosphoethanolamine (POPE) and Cholesterol (Chol): POPC, POPC:Chol (3:1), POPC:POPE (3:1), and POPC:POPE:Chol (2:1:1). The POPC:POPE:Chol (2:1:1) membrane was chosen to mimic the plasma membrane while the others were used to investigate the specific role of PC, PE and Chol lipids. Only POPC:POPE:Chol (2:1:1) membrane was considered for *h*OAT IF conformation as a comparative model. All systems were solvated in water and neutralized with 154 mM NaCl ions to match physiological conditions.

### 2.3. MD simulation setup

Amber FF14SB^42^, Lipids17^43^ and TIP3P^44^ forcefields were used to model protein, lipids and water, respectively. TIP3P-compatible parameters of Na^+^ and Cl^-^ counterions were obtained from Joung and Cheatham^45,46^.

MD simulations were carried out with the Amber18 package^47^ using both CPU and GPU codes for minimization and equilibration, while MD production was performed exclusively on GPU code^48^. Periodic boundary conditions were applied. Non-bonded interactions were explicitly described within a cut-off distance of 10 Å using electrostatic and Lennard-Jones potentials. Long-distance electrostatic interactions were treated using the Particle Mesh Ewald (PME) method^49^. SHAKE algorithm was used to fix bonds involving hydrogen atoms allowing to set the integration time to 2 fs. Production temperature was set at 310K and maintained using a Langevin thermostat^50^. Constant pressure boundary conditions were initially maintained under semi-isotropic conditions using Berendsen barostat^51^.

All systems were initially equilibrated by first minimizing all atomic positions. Then, water molecules were smoothly thermalized from 0 to 100K during 200 ps under (*N,V,T*) conditions. Additional system thermalization up to 310K was then carried out under semi-isotropic (*N,P,T*) conditions for which pressure control was ensured using Berendsen barostat. System boxes were then equilibrated during 5.5 ns. System details (number of atoms and box sizes) are reported in supporting information (Supplementary Table S3). Three independent replicas per lipid bilayer membrane were performed with up to 2 μs (for OF model) and 1 μs (for IF model) MD simulation each, leading to a total of *ca*. 27 μs. Trajectory snapshots were saved every 10 ps.

### 2.4. Analysis

#### Structural analyses

Given the high-confidence model provided by AF2^27^, the reliability of the present OF model folding was evaluated on the secondary structure and the global MFS folding obtained^3^. Structural analyses were performed using the PyTRAJ and CPPTRAJ AMBER modules^52^, VMD^53^ and in-house python scripts. Analyses were performed on equilibrated 1.5 μs long trajectories according to the evolution of time-dependent backbone root-mean squared deviations (Supplementary Figures S2 and S3). z-Dependent pore radius were calculated using the Hole program^54^. 500 snapshots were considered for each trajectory. Interhelical and interdomain interactions were monitored focusing on contacts (< 4 Å) and H-bond interactions. The latter was considered using distance and angle cutoffs set at 3.0 Å and 135°. The minimum fraction threshold was set at 0.1 given the known uncertainties for side chain rotameric states in protein threading techniques. The dynamic cross correlation matrices were calculated over the OF and IF *h*OAT1 MD trajectories in POPC:POPE:Chol (2:1:1) separately, considering only the MFS core.

#### Principal Component Analysis (PCA)

In order to confirm the OF state of the hereby proposed *h*OAT1 model, trajectories were projected on the MFS conformational space obtained from experimentally resolved MFS structures. This conformational space was obtained by performing PCA over a structural data set consisting of MFS proteins available in the Protein Data Bank^3,55^. The MFS dataset including all alternating access states, *i.e*., OF, OF^occ^, IF, IF^occ^ states, are listed in Supplementary Table S4. PCA was achieved by only considering Cα of the MFS twelve transmembrane helices (TMH, see Supplementary Table S5 for transporter MFS core definitions). Dimensionality reduction by PCA points to the main sources of structural variability in the MFS dataset, which thus allows distinguishing IF and OF states as recently proposed^3,55^. Besides, to monitor the OF subspace sampled during MD simulations, a second PCA was also carried out on the MFS backbone using *h*OAT1 OF trajectories. Every membrane was considered.

#### Clustering

Clustering was performed to identify the different subspaces sampled during MD trajectories. Clustering was achieved using the InfleCS approach which take advantage of Gaussian Mixture Models (GMM) to provide insights into the free energy surface^56^. Clustering was carried out by focusing on the first principal component obtained from PCA performed on all trajectories as well as the extracellular distances between TMH1 and TMH7 of *h*OAT1 OF model (residues 212 – 219 and 447 – 454, respectively). Clustering was achieved by using a grid size of 80×80, 5 iterations and from 2 to 16 gaussian functions for GMM. For further details about this method, see Ref.^56^.

## 3. Results and Discussion

### 3.1. Structural patterns of *h*OAT1

#### 3.1.1. Topological overview of the *h*OAT1 model

In agreement with previous studies^24,25^ as well as AF2 prediction^27^, the present MD-refined model of *h*OAT1 adopted the MFS fold. Despite the low sequence similarity within the MFS superfamily, they share a common architecture. MFS fold mostly consists of 12 transmembrane helices (TMHs) divided into two bundles of 6 TMHs each. The so-called N- and C-bundles comprise TMH1-6 and TMH7-12, respectively (see Figure 1a&b)^57^. As expected for an MFS-fold transporter^3,55,58,59^, N- and C-domains exhibited pseudo-symmetry perpendicularly to the plane of the membrane (Figure 1a). The present model revealed at least 6 intracellular helices (ICHs, Figure 1a&b), as observed in the AF2 model^27^ and other experimentally resolved mammalian MFS transporters (*e.g*., GLUT1/*SLC2A1*, GLUT3/*SLC2A3*, GLUT5/*SLCA5*)^37,60,61^. These ICHs are in close contact with TMHs for which local details are discussed in section 3.2. Finally, *h*OAT1 topology suggested a long extracellular loop (ECL1) made of *ca*. 90 amino acids (40-126) between TMH1 and 2. The secondary structure of ECL1 appeared more disordered in the present model than in AF2, leading to lower confidence for ECL1 than for MFS core. However, this is not expected to strongly affect the MFS core structure discussed in the present study. Furthermore, the glycosylation of known sites in ECL1 is not required for the transporter function^14,28^.

**Figure 1.**
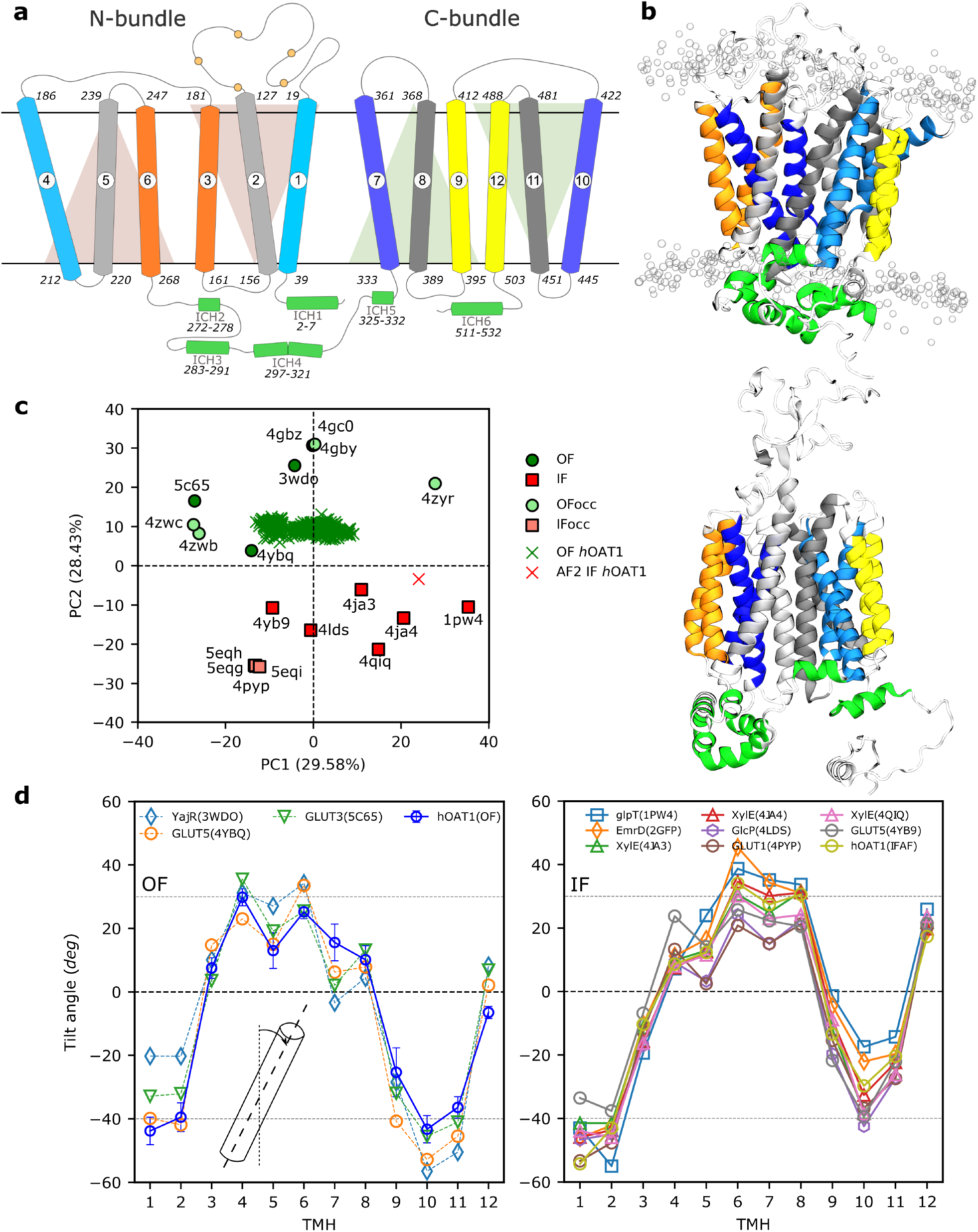
Overview of the *h*OAT1 transporter. (a) The topology scheme shows *h*OAT1 adopting the canonical MFS fold that consists of 12 transmembrane helices (TMH) divided into N- and C-bundles, connected by an intracellular loop rich in intracellular helices (ICHs). Each bundle is constructed of 3-TMH inverted segments. TMH1 and TMH2 are connected by a long extracellular loop possessing 5 glycosylation sites (Arg39, Arg56, Arg92, Arg97, Arg113). The so-called A-, B- and C-helices are depicted in blueish, grayish, and yellowish, respectively. (b) 3D model of *h*OAT1 obtained from MD simulation and AlphaFold2 prediction in OF (top) and IF (bottom) conformational states, respectively. (c) *h*OAT1 projected onto the conformational space obtained via PCA of resolved MFS transporters in OF, OF^occ^, IF, IF^occ^ conformations. (d) Tilt angle profile of MFS transporters in OF (left) and IF (right) conformations. The TMH tilt angle profile for the *h*OAT1 OF model was averaged over MD simulations considering all replicas.

#### 3.1.2. hOAT1 model adopts OF state conformation according to MFS conformational space

MFS alternating access is expected to follow the rocker-switch mechanism^3^ in which N- and C-bundles rearrange between OF and IF states. This large-scale conformational change along the transport cycle was also shown to affect intra-bundle TMH arrangements^3^. Overall, the MFS tertiary structure is modified along the transport cycle by rocking N- and C-bundles to alternatingly expose substrates to both sides of the lipid bilayer membrane (see Ref.^3^ for further details). Visual comparison between the MD-refined *h*OAT1 model and AF2 prediction suggested two distinct states. The present model adopted a “V”-shape conformation, while AF2 clearly exhibited a “A”-shape (Figure 1b&c) as proposed for the IF conformation of resolved MFS transporters^24,25^. This was confirmed by projecting MD trajectories and the AF2 structure onto the MFS conformational space obtained by PCA (see Figure 1c and Supplementary Figure S4). Besides, the OF state was also confirmed by exhibiting significantly larger (respectively smaller) Met358-Ser139 (Gly446 - Val211) distances with respect to the previous IF model obtained by Tsigelny *et al*.^25^. These distances were suggested to picture opening at either the extra- or intra-cellular sides, respectively (see Supplementary Figure S5).

Building upon the concept of typical structural features for MFS proteins, tilt angles between TMHs and the lipid bilayer axis normal were monitored along MD simulations (see Supplementary Figure S6) and compared with experimentally resolved MFS transporters. Tilt angle profiles were averaged over MD trajectories, exhibiting good agreement with the profiles obtained with experimentally resolved OF state MFS transporters (see Figure 1d). The present OF *h*OAT1 model exhibited the well-known 3-TMH repeated segment fold observed in MFS proteins. Within each bundle, two 3-TMH segments are related by approximately 180° rotation around the lipid bilayer normal^3,57,58^. This leads to three sets of TMHs, namely A-, B- and C-helices (see Figure 1 and Supplementary Figure S7) for which different functional roles were suggested^59^. The dynamic interplay of interactions between helices is an imperative part of alternation between OF and IF states, including the existence of intermediate occluded states^59^. The averaged contact maps of OF and IF states were also compared along MD simulations, focusing on the MFS core (see Supplementary Figure S8). Overall, intra-bundle contacts were conserved between the two states. This is in agreement with the suggested rocker-switch mechanism^3^ for MFS transport cycle. Furthermore, in line with the large-scale conformational changes occurring during the transport cycle, the strong interdependency between TMHs was suggested from MFS core dynamic cross correlation matrix; showing significant motion correlations between TMHs along simulations (see Supplementary Figure S9).

#### 3.1.3. Structural arrangement of TMHs in *h*OAT1 OF model

It is important to note that A- and B-helices were suggested to act by pairs across N- and C-bundles. Therefore, in the present section, particular attention was paid to inter-bundle interactions for A- and B-helices in contrast to C-helices.

The central cavity of *h*OAT1 consists of A-helices, namely TMH1 and 4 for the N-bundle and TMH7 and 10 for the C-bundle (see Figure 1 and Supplementary Figure S7). These helices play a role in substrate binding and release events in the OF and IF states, respectively^59,61,62^. They interact by pairs across bundles *i.e*., TMH1 with TMH7 and TMH4 with TMH10. Key non-covalent interactions between TMH1 and 7 occur on the extracellular site. z-Dependent cavity pore radii exhibited the expected OF pattern, *i.e*., greater opening in the extracellular side than in the intracellular one (see Supplementary Figure S10). However, large standard deviations suggest the existence of OF occluded structures during MD simulations. This was confirmed by monitoring (i) the extracellular distance between TMH1 and TMH7 (see Supplementary Figure S11) and (ii) performing PCA considering all lipid bilayers (see Supplementary Figures S12 and S13). The first two principal components (PC) were assigned to the opening and closing of the extracellular gate, for which TMH contributions are reported in Supplementary Figure S14. PC1 and the extracellular distance between TMH1 and TMH7 were used to picture the OF subspace sampled during MD simulations by means of the InfleCS method^56^. It is important to note that the free energy barriers obtained from InfleCS should be carefully considered given the low sampling of intermediate regions. However, three main state populations were clearly identified, namely extracellular open, intermediate, and closed conformations (see Figure 2). This confirms the central role of the TMH1/TMH7 pair which is expected to be involved in extracellular gating event prior to the occlusion of the extracellular gate along transport cycle. MD simulations revealed the following interacting residues in *h*OAT1: Asn35, Thr36, Asn39, Phe40 for TMH1 and Tyr353, Tyr354, Leu356, Val357 for TMH7. MD simulations also revealed kinking of A-helices, in agreement with previous studies^57,59^. Investigations of TMH helicities in OF *h*OAT1 model exhibited discontinuity in TMH1 and 10 (see Supplementary Table S6) leading to elbow-shape TMHs. A-helix discontinuities were used to picture the structural adaptability of MFS core along the OF-to-IF transition and *vice versa*^38,57,59^.

**Figure 2.**
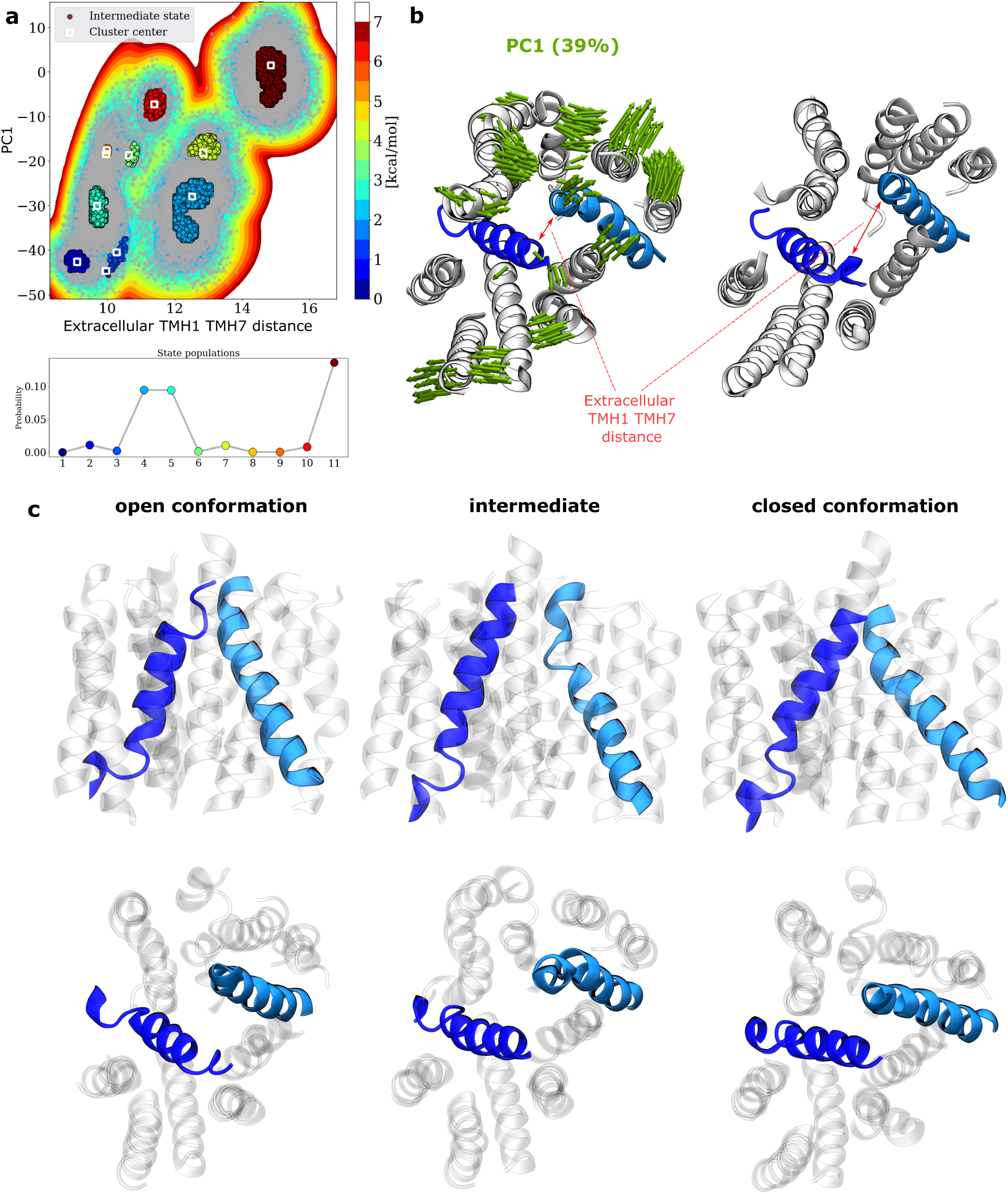
Conformational sampling of extracellular gating events of *h*OAT1 OF model. (a) Insights into the free energy surface (top) sampled during MD simulations according to PC1 and extracellular distance between TMH1 and TMH7 as well as cluster probabilities (bottom). Porcupine plot (left) obtained from PCA performed considering *h*OAT1 model embedded in all lipid bilayer membranes and corresponding evolution of extracellular distance between TMH1 and TMH7. (c) Representative snapshots of the three main clusters in which TMH1 and TMH7 are highlighted to feature occlusion states (side and top are respectively shown on top and bottom panels.

It provides flexibility allowing side chains of gating residues to interact within paired A-helices. Structural analyses performed on OF *h*OAT1 model enabled the identification of dispersive, electrostatic, and H-bond interactions between the aforementioned residues involved in the so-called “gating” events. MD simulations and the AF2 model support the key role of Tyr354/Tyr353 placed on the “elbow” point of TMH7 for gating as it was shown for conserved tyrosine in sugar porters (*e.g*., conserved Tyr292 and Tyr293 in *h*GLUT1, Tyr290 and Tyr291 in *h*GLUT3)^3^. MD simulations stressed out that interactions between A-helices are highly dynamic as pictured by H-bond network analysis (see Supplementary Tables S7 and S8), especially for extracellular occlusion event in OF *h*OAT1. Our simulations suggest that the interchange between the OF open and occluded states can dynamically occur even in the absence of substrate, owing to local flexibility and thermal fluctuation^55^ as pictured by the aforementioned clusters (see Figure 2 and Supplementary Figure S7).

B-helices (TMHs 2, 5, 8 and 11, see Figure 1a and Supplementary Figure S7) are expected to play a role in maintaining the interface between the N- and C-bundles^59^. As shown for A-helices, B-helices might be considered as pairs, *i.e*., TMH2/TMH11 and TMH5/TMH8. Therefore, particular attention was paid to non-covalent interactions between bundles. The present *h*OAT1 model is in agreement with these findings as pictured for instance by strong H-bond interactions between TMH5 and TMH8, which are maintained for more than 80% of the time during MD simulations (see Supplementary Tables S7 and S8). The most frequent residues involved in TMH2/TMH11 and TMH5/TMH8 H-bond networks are reported in Supplementary Table S8. Besides, in line with previous experimental observations, OF *h*OAT1 B-helices are likely to participate in substrate binding and translocation along transport cycle thanks to: (i) their “banana-shape” bending (see *e.g*., TMH2 in Supplementary Figures S7 and S13)^38^; and (ii) their residues exposed at the substrate cavity (*e.g*., Arg466, Ser469, Arg131, Arg134). In OF state conformation, TMH5/TMH8 interactions are preserved all along the helices. Bending of B-helices displays a different profile for AF2 IF with respect to OF *h*OAT1 conformation. The helices differ in the curvature at the helical ends, and the most pronounced variation between states was found for TMH11 and TMH8 (Supplementary Figure S7). This suggests that large-scale conformational changes along the *h*OAT1 transport cycle are asymmetric. The C-bundle is likely to be more flexible, in line with previous findings regarding other MFS proteins such as LeuT, *h*GLUT3, and *h*GLUT5^3,38,61,63^. This hypothesis was strengthened by both PCA and TMH tilt angle profiles obtained with the OF *h*OAT1 model, which showed larger flexibility for the C-than for the N-bundle (see Supplementary Figures S4 and S13). This is also the case for AF2 (see Supplementary Figure S3).

Finally, the C-helices (TMH3, TMH6, TMH9 and TMH12, see Figure 1a and Supplementary Figure S7) are located out of the central *h*OAT1 core. In contrast to A- and B-helices, C-helices stand by each other in each bundle, *i.e*., TMH3/TMH6 and TMH9/TMH12 for the N- and C-bundles, respectively. They support the structure integrity of *h*OAT1 by interacting with the lipid bilayer. In the present OF model, inter-helical interactions between TMH3 and TMH6 are mostly located at the intracellular side. This is not the case for TMH9 and TMH12 which exhibit contacts over the whole helices. Interestingly, the opposite trend seems to occur with the AF2 IF *h*OAT1 model: TMH9/TMH12 exhibit less contact than TMH3/TMH6, likely due to a conformational change along the transport cycle.

### 3.2. The importance of MFS conserved sequences on the “charge-relay” system of *h*OAT1

#### 3.2.1 MFS conserved motifs as central components of the charge-relay system

*h*OAT1 shares with other MFS transporters conserved sequences across species, which were shown to act as “molecular switches” during the transport cycle by *e.g*., triggering large-scale conformational changes^31,64,65^. The so-called MFS signature motifs are located at the intracellular interface, *i.e*., in intracellular loops (ICLs) and TMHs as observed in other MFS transporters^57,59,64,66,67^. These motifs are duplicated in the N- and C-bundles. MFS signature motifs identified in *h*OAT1 may slightly differ in terms of sequence between the two bundles (see Table 1 and Figure 3), as well as with other MFS proteins^24,68^. The so-called A-motifs^3,64,65^ are located in the ICLs between TMH2 and TMH3 in the N-bundle, and between TMH8 and TMH9 in the C-bundle. The N-bundle A-motif matches with the canonical sequence, *i.e*., G[X3]D[R/K]XGR[R/K]. The C-bundle A-motif sequence is shorter, whilst the final LGRR pattern is conserved (Table 1 and Figure 3). The E[X_6_]R sequence (also known as ELYPT^66,68^ for the N-bundle) is observed in the ICLs connecting TMH4 and TMH5 in the N-bundle and TMH10 and TMH11 in the C-bundle. The PETL motif is located in the C-terminal domain, after TMH12. Finally, the conserved [P/X]ESXRW[L/X] sequence^66,68^ is also observed in *h*OAT1 after TMH6, in the intracellular domain connecting the N- and C-bundles. In spite of significantly different primary sequences, OF and IF *h*OAT1 structural models support that PETL and [P/X]ESXRW[L/X] motifs are expected to behave similarly in the C- and N-bundles, respectively^3^. Thereby, it must be stressed that, for sake of readability, both motifs will be referred to as PETL.

**Table 1.**
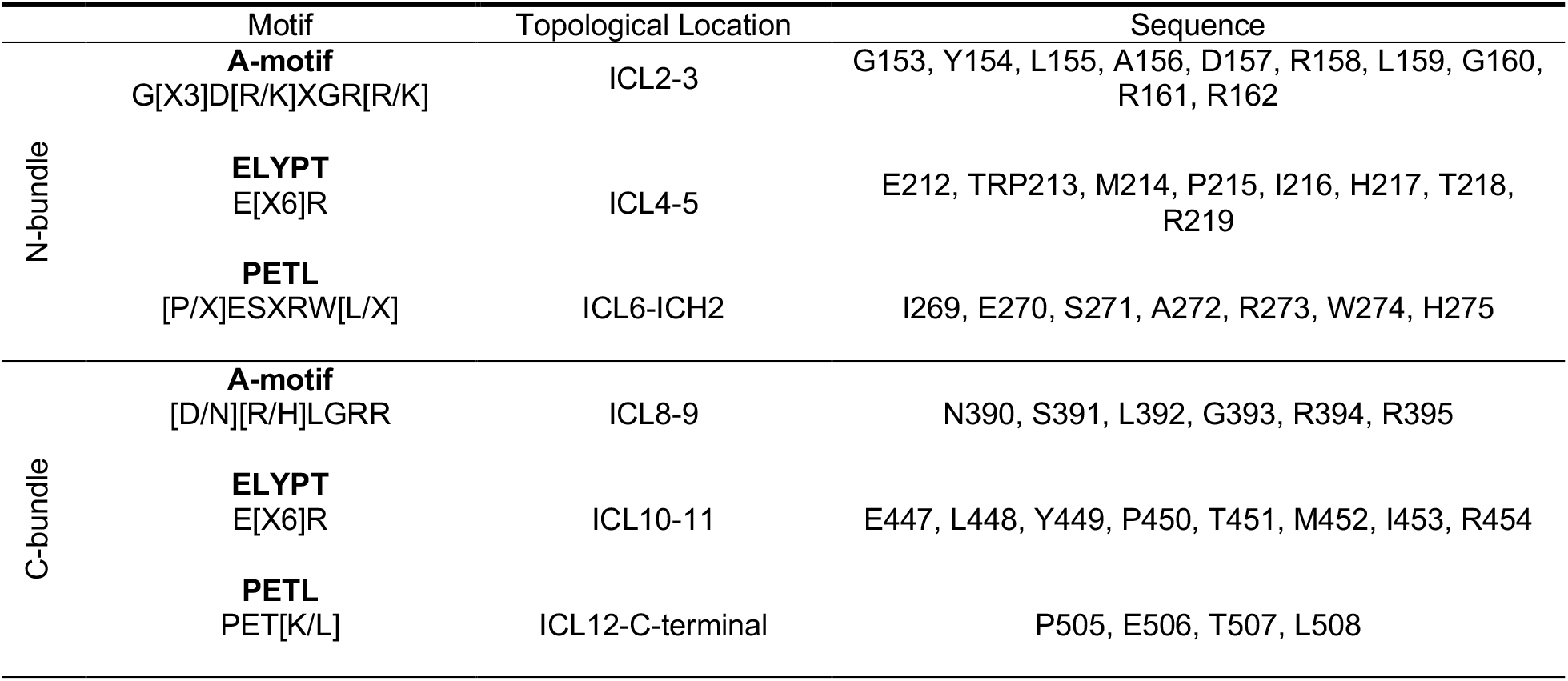
The description of MFS signature intracellular motifs found in *h*OAT1 divided into N- and C-bundles showing the pseudosymmetry of the transporter.

|  | Motif | Topological Location | Sequence |
| --- | --- | --- | --- |
| N-bundle | **A-motif**<br>G[X3]D[R/K]XGR[R/K] | ICL2-3 | G153, Y154, L155, A156, D157, R158, L159, G160, R161, R162 |
|  | **ELYPT**<br>E[X6]R | ICL4-5 | E212, TRP213, M214, P215, I216, H217, T218, R219 |
|  | **PETL**<br>[P/X]ESXRW[L/X] | ICL6-ICH2 | I269, E270, S271, A272, R273, W274, H275 |
| C-bundle | **A-motif**<br>[D/N][R/H]LGRR | ICL8-9 | N390, S391, L392, G393, R394, R395 |
|  | **ELYPT**<br>E[X6]R | ICL10-11 | E447, L448, Y449, P450, T451, M452, I453, R454 |
|  | **PETL**<br>PET[K/L] | ICL12-C-terminal | P505, E506, T507, L508 |

**Figure 3.**
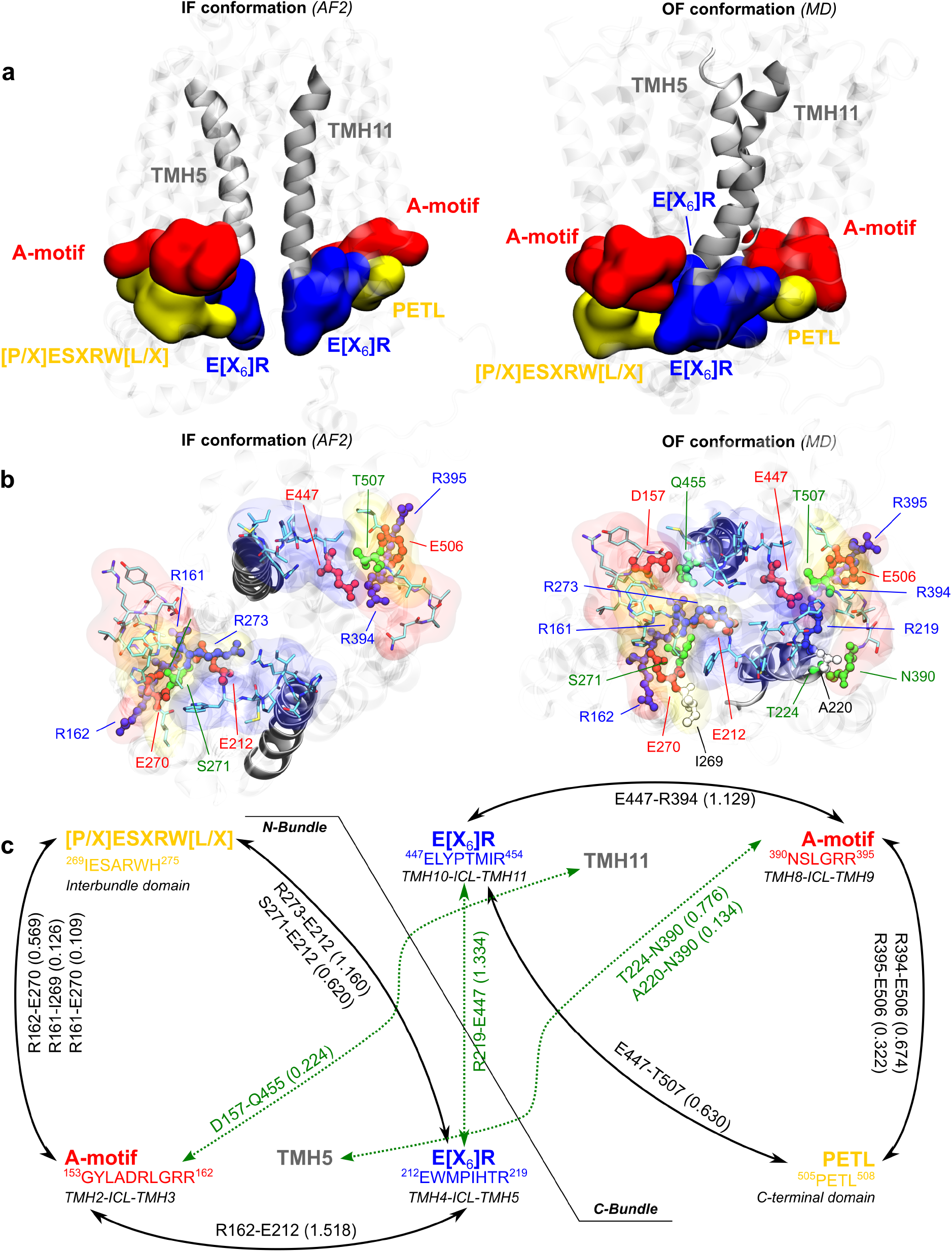
Intracellular motifs conserved among MFS. (a) Charge-relay system of *h*OAT1 as a triad made of A-motif, [P/X]ESXRW[L/X] / PETL and E[X_6_]R symmetrically in the N- and C-bundles visualized in IF (left) and OF (right) conformations. (b) Intracellular view of the charge-relay system with highlighted residues involved in interactions. (c) The map of each motif interactions emphasizing the symmetry in bundles. The communication within motifs is demonstrated by the strength of hydrogen bonds. Green dotted lines represent the missing interactions in the IF model, crucial for conformational changes. It must be stressed that values above 1.0 highlight salt-bridges in which more than one H-bond is possible (*e.g*., between arginine and glutamate/aspartate residues).

MFS signature motifs are rich in charged and polar amino acids (mostly arginine, aspartate, and glutamate) leading to strong H-bond and salt-bridge networks. MD simulations showed that the so-called “charge-relay system”^64,65^ is highly dynamic since salt-bridges and H-bonds can be exchanged along the simulation. The IF AF2 and MD-refined OF models were then used to identify shared patterns and conformation-dependent rearrangements.

The “charge-relay system” can be divided into two building blocks in the N- and C-bundles, each made of A-motif, E[X6]R, PETL motifs (Figure 3a). These motifs share a similar structural arrangement regardless of the conformational state. H-bond analyses highlighted the central role of the last two arginine residues of A-motifs in maintaining the supramolecular arrangement with the other two motifs (see Figure 3b&c). N-bundle Arg161 and Arg162 interact with Glu212 and Glu270 from the E[X_6_]R_N-bundle_ and PETL_N-bundle_ motifs, respectively. Likewise, in C-bundle, Arg394 and Arg395 interact with Glu447 and Glu506, respectively in the E[X_6_]R_C-bundle_ and PETL_C-bundle_ motifs. It is worth mentioning that our findings are supported by a directed site-mutagenesis experiment where the mutation of Glu506 led to complete inactivation of *h*OAT1 transport^29^. Glu212 and Glu447 in the N- and C-bundle E[X_6_]R motifs also interact with PETL motifs. Using MD simulations on the OF *h*OAT1 model, H-bond fractions over time were also calculated to measure the strength of the local H-bond network in each triad (Figure 3c).

Salt-bridges between A- and E[X_6_]R motifs exhibit highly conserved interactions for Arg162-Glu212 and Arg394-Glu447; time fractions respectively being above 1.0 along MD simulations (see Figure 3c and Supplementary Table S9). Interestingly, contact analysis of static AF2 IF conformation suggested similar H-bond pattern (see Figure 3b). This was confirmed by monitoring H-bond during MD simulations performed on the IF *h*OAT1 model (see Supplementary Table S9). Similar H-bond network as OF conformation was observed, supporting the existence of the conformation-independent motif arrangement within each bundle.

Interestingly, interactions between motifs across bundles differ significantly in IF and OF models. The AF2 IF *h*OAT1 model does not exhibit non-covalent interactions, nor even contacts, between motifs of N- and C-bundles (see Figure 3a&b). In contrast, *h*OAT1 OF model exhibits strong H-bond and salt-bridge networks (see Supplementary Table S9). The supramolecular arrangement of OF *h*OAT1 relies on the interactions between the two E[X_6_]R motifs, as pictured by the strong salt bridge between Arg219 and Glu447 (H-bond fraction of 1.334). In agreement with previous observations on MFS proteins^3,37,55,59,65^, MD simulations show that cross-bundle interactions also involved the intracellular side of TMHs with MFS signature motifs, but to a lesser extent. For example, H-bonds were observed between N-bundle A-motif and TMH11 (Asp157-Gln455, fraction = 0.224) or between C-bundle A-motif and TMH5 (Thr224 or Ala220 with Asn390, with fractions of 0.776 and 0.134, respectively). H-bond interactions were also monitored on IF *h*OAT1 trajectories; revealing the absence of inter-bundle H-bond (see Supplementary Table S9) leading to a greater distance between E[X_6_]R motifs (*ca*. 12 and 17Å, respectively for OF and IF conformations, see Supplementary Figure S15).

This suggests that A-motifs might be involved in locking intracellular gate, which in turn maintain the OF conformation. This is in agreement with previous observations on resolved MFS proteins adopting OF confirmations, such as YajR and GLUT1^3,37,55,59,65^. For example, high-throughput single directed mutagenesis performed on glycine and aspartic acid in the first and the fifth position of the YajR MFS transporter A-motif showed conformational transition from OF-to-IF, while other single point mutations only destabilized the protein^31,55,62,65^. Likewise, the E[X_6_]R motif may play an important role in local arrangement of TMHs across bundles in OF conformation. Interestingly, structural analyses revealed only interactions of the N-bundle E[X_6_]R motif with TMH11 through the H-bond interaction between Glu212 and Gln455 (fraction=0.464). Comparatively it was shown for the YajR transporter, where the interactions between TMH2 and TMH11 would be essential for the OF conformation^65^. However, no interaction was observed between C-bundle E[X_6_]R and TMH5 in spite of the pseudosymmetry of the MFS transporter. This may be due to the resolution of the present OF model. Besides, it may also suggest an asymmetrical behavior in *h*OAT1 between the N- and C-bundles, which requires further investigations.

Despite the lower confidence of our model regarding the resolution of intracellular loops and helices in the OF state, the MD simulations as well as the comparison with AF2 structure provided hints regarding the cytoplasmic arrangement. As observed for resolved GLUTs^38,55,61,62^, both models suggest that intracellular helices (ICHs) are in contact with the MFS signature motifs. In the AF2 IF model, ICHs are separated between the N- and C-bundles, while the OF model suggest contacts between ICHs as well as with intracellular loops. These interactions are expected to play a key role along the transport cycle. In case of sugar porters (*e.g*., GLUT1 or GLUT3), ICHs and TMHs were proposed to lock the transporter in the OF conformation, precluding the exposure of the intracellular gate to the environment^3,55,61^.

Altogether, present MD simulations findings line up with the putative role of tightly arranged intracellular interactions engaging the ICHs that are likely involved in substrate access to the intracellular gate. It is consistent with previous hypotheses that the intracellular interactions of *h*OAT1 are also prompt to stabilize the OF conformation. Therefore, the eventual breakage of these interactions may be directly involved in the conformational change along the transport cycle^12,59,61,62,64,65^.

### 3.3. The impact of membrane lipid components

#### 3.3.1. On the interplay between lipid composition and the *h*OAT1 conformational space

MFS transporter structures and functions were both computationally and experimentally shown to strongly depend on membrane composition^3,30–32,59^. This is particularly true for membranes made of PC and mixtures of PE phospholipids, which showed different behaviors in term of non-covalent interactions with membrane proteins^30–32,69,70^. In the present study, MD simulations were used to provide insights in protein-membrane interactions. The OF *h*OAT1 model was embedded in various lipid bilayers, *i.e*., POPC:POPE:Chol (2:1:1), POPC:Chol (3:1), POPC:POPE (3:1), and POPC, the first one presenting the closest amounts of PE lipids and cholesterol to actual cell membranes^71^. Although the membranes used in the present study do not comprehensively account for the whole complexity of cell membranes in terms of composition and asymmetry, they are expected to faithfully catch the main features of membrane-protein interactions for the most abundant lipids, *i.e*., PC, PE and cholesterol.

In order to assess the overall lipid composition-structure relationship, trajectories obtained from MD simulations in different lipid bilayer membranes were all projected onto the MFS conformational space obtained using PCA on the resolved structure. Regardless of the membrane composition, all systems conserved the expected OF conformation along simulations as shown by PCA projection as well as TMH tilt angle profiles (see Supplementary Figures S16 and S17). However, PCA projections revealed that protein dynamics and conformational space are slightly different in pure POPC and binary lipid bilayer (*i.e*., POPC:POPE and POPC:Chol), as compared to the POPC:POPE:Chol (2:1:1) lipid bilayer membrane. Trajectories were also projected on the OF subspace obtained from the PCA calculated using all lipid bilayer membrane (see Supplementary Figure S12). PCA projection supports a differential behavior according to lipid bilayer composition (see Supplementary Figure S18). In absence of POPE and/or cholesterol, replicas sampled different subspaces with no overlap between them. This suggest that PE lipids and cholesterol play a central role in *h*OAT1 dynamics. MD simulations also suggested that lipid composition is also expected to slightly modulate gating events as pictured by the extracellular distances between TMH1-TMH7 (see Supplementary Figure S11). Although no clear conclusion in terms of function can be drawn from these results, differential protein dynamics according to lipid bilayer composition are in agreement with previous observations^30,31^ stressing the importance of protein-lipid non-covalent interactions.

Both intracellular and extracellular openings were monitored by respectively measuring Met358-Ser139 and Val211-Gly446 distances in *apo h*OAT1 simulations (Supplementary Figure S5). Given that *h*OAT1 adopts the OF conformation, intracellular distances were expected to exhibit low variability owing to the structural intracellular arrangement maintained by the charge-relay system (see Supplementary Figure S11). Interestingly, intracellular distances exhibit slightly higher variability in a PE-free than in a PE-based lipid bilayer membrane. This may picture a looser packing of intracellular loops which may in turn modulate *h*OAT1 function.

#### 3.3.2. Non-covalent interactions between lipid components and the *h*OAT1 transporter

The ability of lipid bilayer membranes to form H-bond networks is expected to contribute to protein stability and dynamics. To probe protein-lipid interactions, H-bond networks were monitored along trajectories. It is important to note that protein-lipid H-bond interactions are highly dynamic^30,31^ leading to lipid-lipid exchange along the trajectories. The strongest network was observed in the POPC:POPE:Chol (2:1:1) membrane (Figure 4a). This is explained by the higher H-bond donor feature of PE polar heads, with their ammonium N-atom, than of PC polar heads. Interestingly, in the absence of PE lipids, PC contribution to H-bond networks increases. It is worth mentioning that many less H-bonds were observed in cholesterol-protein interactions, owing to the single OH group of cholesterol. However, the presence of cholesterol in lipid bilayer membranes tends to favor PC- and PE-protein H-bond interactions. Cholesterol is known to modulate lipid dynamics by *e.g*., increasing lipid order and dynamics in fluid lipid bilayer membranes^59^. Furthermore, the presence of cholesterol in artificial membranes is associated with local low and high lipid packing through lipid-lipid interactions^3,59^. Therefore, the presence of cholesterol is expected to decrease PC and PE lipid dynamics, which in turn increases presential lifetime of surrounding lipids. Given the high dynamic feature of lipid-protein interactions, distributions of surrounding lipids were also calculated focusing on cholesterol and PE polar heads.

**Figure 4.**
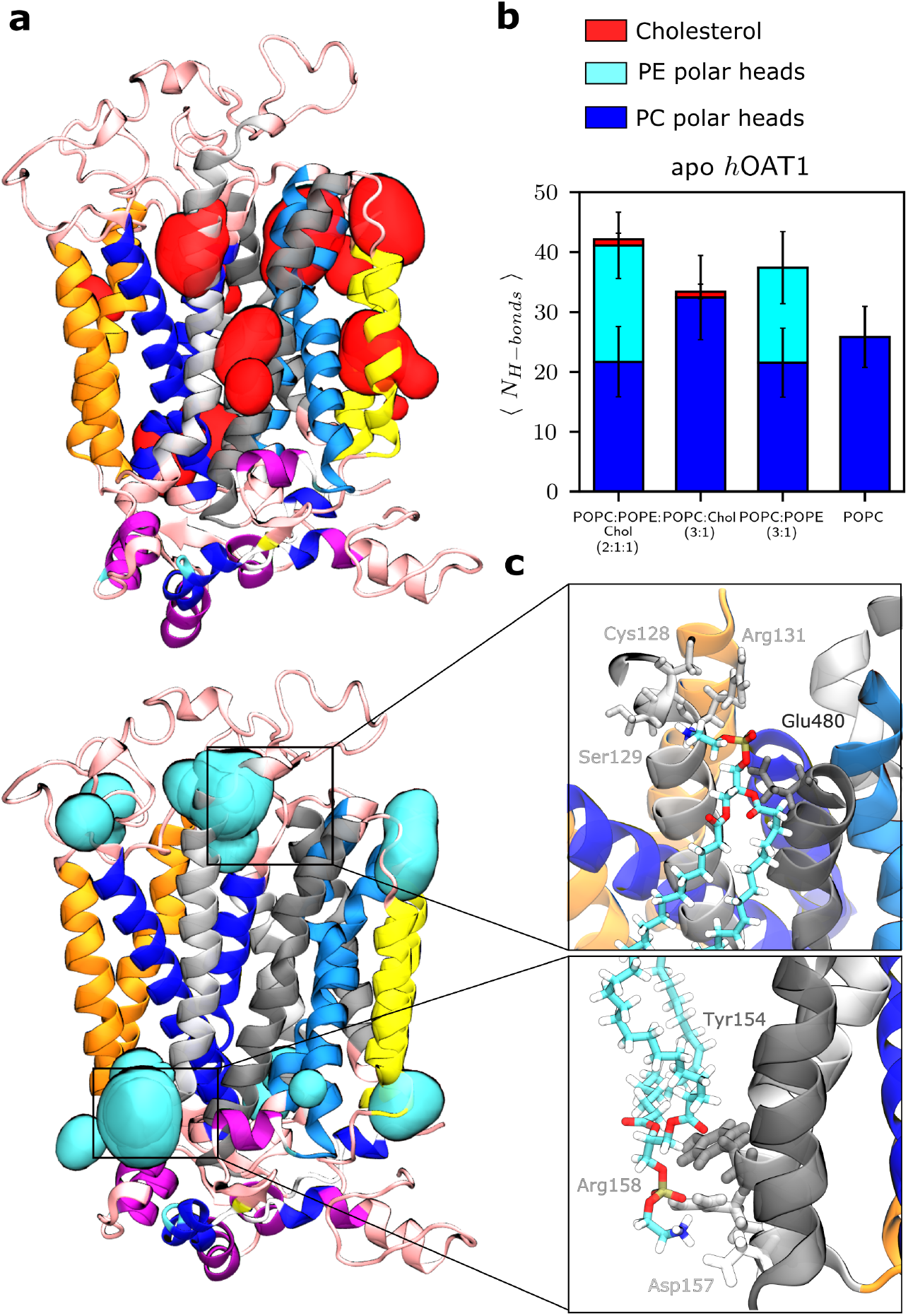
Impact of the membrane lipid components. (a) Hot spots for lipid-protein interactions appearing over 80% of simulations. (b) Number of hydrogen bonds between lipid polar heads and *h*OAT1 for each membrane. (c) Close-up frame points for specific interactions: PE polar heads disrupt salt-bridges between gating residues placed on the extracellular ends of TMH2 and TMH11 (top); PE polar heads interacting with Tyr154, Asp157 and Arg158 by the A-motif (bottom).

The calculated protein-lipid density maps suggested several hotspots for B- and C-helices where *h*OAT1 residues are preferentially in contact with either PE polar heads or cholesterol for more than 80% of the simulations (see Figure 4b and Supplementary Figure S19). It suggests that specific lipid binding sites exist, in agreement with observations made for other MFS transporters^30–32^. Present analyses exhibited that cholesterol might have strong binding in the TMH1 region of the inner leaflet, as well as in the TMH8 and TMH10 regions of the upper leaflet. Other hot spots were observed between TMH9 and TM12. Regarding PE lipids, a binding site was observed involving residues located on the extracellular site of TMH2 and TMH11, in agreement with observations made for XylE and LacY transporters^31^. Direct interactions between PE and TMH2 and TMH11 were suggested to modulate the conformational state dynamics^30–32,69,70^. For instance, in GLUT transporters, PE lipids were shown to compete over the salt-bridges between N- and C-bundles^31^. In the present OF *h*OAT1 model, PE lipids disrupt the salt bridge between Glu480 and Arg131/Arg138 which ultimately may stabilize the OF state (Figure 4). Regarding protein-lipid interactions on the intracellular side, conformational changes along the transport cycle were reported to be facilitated by lipids through lipid-A-motif interactions^31^.

MD simulations proposed that PE lipids preferentially interact with Tyr154, Asp157 and Arg158 which are involved in the charge-relay system. This event was not observed with PC lipids. Even though the present results should be considered carefully, they stress out the central role of PE lipids in *h*OAT1 dynamics and function in agreement with observations made for other MFS transporters. For instance, PE lipids were shown to act as a chaperon facilitating the folding of LacY transporter^72^. Function-wise, lipids were shown to disrupt key salt-bridges which in turn may favor conformational changes along the transport cycle^3^. For example, LacY transporter activity was increased in the presence of PE lipids^73,74^. The same was shown for the xylose (XylE) and Glycerol-3-phosphate (GlpT) transporters, the conformational states of which were also stabilized by PE lipids^31^. Several transporters possess a cholesterol binding site with a distinct role^59^. In GLUT transporters, the presence of cholesterol has been found to stabilize the protein and potentially promote oligomerization^75,76^. Besides, the presence of PE lipids is known to increase membrane fluidity and thus contribute to lipid packing defects^59^. PE components would facilitate the transport cycle by direct interactions with key residues of the transporter^31^.

## 4. Conclusion

We propose a novel, full molecular model of the human *SLC22A6*/OAT1 transporter in the outward-facing conformation. The present model was thoroughly compared with the recently proposed IF *h*OAT1 model obtained using AF2 and validated by using the conformational space from experimentally resolved MFS transporters. Particular attention was paid to the transmembrane domain for which TMH arrangements are consistent with the OF conformation and literature reports. The role of *h*OAT1 intracellular charge-relay system was investigated, highlighting key residues involved in salt bridges. Comparison with the AF2 IF *h*OAT1 model suggests the existence of two local intracellular arrangements in which conserved motifs may lock *h*OAT1 in the OF conformation. The conformational change is likely facilitated by specific interactions of PE lipid components with gating and motif residues confirming the dependency of MFS proteins on the composition of lipid bilayer. The present model can be used for further investigation of drug(-drug) interactions (inhibitory studies) by providing atomic pictures and binding affinities for given drugs.

Finally, the present model should help to better understand *h*OAT1 function at the molecular level, pending experimental resolution by means of *e.g*., cryo-EM techniques. This model should help rationalize known polymorphism or rare mutation by *e.g*., simply replacing amino acid of importance and achieving MD relaxation. It can also be used to investigate local binding models of small molecules to support substrate/inhibitor competitive experiments.

## Abbreviations

ABC: ATP-Binding Cassette
AF2: AlphaFold 2
*a*KG: *α*-ketoglutarate
Chol: Cholesterol
Cryo-EM: Cryogenic Electron Microscopy
ECL: ExtraCellular Loop
GlpT: Glycerol-3-phosphate Transporter
GLUT: Glucose Transporter
H-bond: Hydrogen bond
ICH: IntraCellular Helix
ICLs: IntraCellular Loops
IF: Inward-Facing
IF^occ^: Inward-Facing occluded
LacY: Lactose permease
LeuT: Leucine Transporter
MD: Molecular Dynamics
MFS: Major Facilitator Superfamily
NaDC3: Na^+^/Dicarboxylate Cotransporter
NKT: New Kidney Transporter
OAT: Organic Anion Transporter
OF: Outward-Facing
OF^occ^: Outward-Facing occluded
PC: PhosphatidylCholine
PCA: Principal Component Analysis
PD: PharmacoDynamics
PE: PhosphatidylEthanolamine
PGx: Pharmacogenetics
PK: PharmacoKinetics
PME: Particle Mesh Ewald
POPC: 1-palmitoyl-2-oleoyl-sn-glycero-3-phosphocholine
POPE: 1-palmitoyl-2-oleoyl-sn-glycero-3-phosphoethanolamine
PTC: Proximal Tubular Cells
SLC: SoLute Carrier
SNP: Single Nucleotide Polymorphisms
TMH: TransMembrane Helix
XylE: Xylose transporter

## 6. Acknowledgments

This work was granted access to the HPC resources of IDRIS under the allocations 2020-A0080711487 and 2021-A0100711487 made by GENCI, using the GPU supercomputer “Jean Zay”. We are grateful to regional supercomputer CALI (“CAlcul en LImousin”) and Xavier Montagutelli for technical support.

## 8. Data availability

The datasets generated during the current study (*e.g*., structures, trajectories, structural parameters, molecular dynamics inputs are available from the corresponding author on reasonable request.

