## Supplementary Information for "Insights into the structure and function of the human organic anion transporter 1 in lipid bilayer membranes"

INSERM U1248 IPPRITT, CBRS, Faculté de Médecine et Pharmacie,

Univ. Limoges,

87000 Limoges

France

### 1. Supplemental Tables

**Supplementary Table S1.** *In vitro* studies of site-directed mutagenesis performed on OAT1.

| Amino acid change | Localization | Substate | Effects | Cell model (*in vitro*) | Reference |
|---|---|---|---|---|---|
| p.Leu30Ala | TMH1 | PAH | Loss of expression | COS-7 | [14] |
| p.Thr36Ala | | | Loss of substrate recognition | | |
| p.Asn39Glu | TMH1 | PAH | Loss of transport activity; no effect of cell surface expression | HeLa | [15] |
| p.Asn(39/56/92/97)Glu | TMH1/ECL | PAH | Loss of transport activity and cell surface expression | | |
| p.Tyr230Ala/Phe | TMH5 | CDF PAH | Loss of membrane protein expression and diminution of transport activity | Xenopus Oocyte | [16] |
| p.Lys431Ala/Tyr | TMH10 | | Diminution of membrane protein expression and transport activity | | |
| p.Phe438Ala/Tyr | TMH10 | | Diminution of membrane protein expression and transport activity | | |
| p.Tyr490Ala<br>p.Leu503Ala<br>p.Leu504Ala | TMH12 | PAH | Loss of membrane protein expression and transport activity | COS-7 and LLC-PK1 | [17] |
| p.Arg466Lys | TMH11 | PAH | Decreased transport activity but no impact on expression | Xenopus Oocyte | [18] |
| p.Ser203Ala | TMH4 | AFV CDF | Decreased transport activity | HEK293 | [19] |
| p.Tyr353Ala<br>p.Tyr354Ala | TMH7 | PAH | Loss of transport activity | COS7 | [20] |
| p.Trp346Ala<br>p.Thr349Ala | | / | Loss of transport activity and of the total expression | | |
| p.Glu506Ala/Gln | ICH6 | PAH | Decreased transport activity | COS-7 | [21] |
| p.Leu512Ala | | | | | |
| p.Cys440Ala | TMH10 | PAH | Decreased transport activity | CHO cell | [22] |
| p.Cys189Ala | TMH4 | | | | |
| p.Cys341Ala | TMH7 | | | | |

| | | | | | |
|---|---|---|---|---|---|
| p.Cys206Ser | TMH4 | | | | |
| p.Cys335Ala | TMH7 | | | | |
| p.Cys379Ala | TMH8 | PAH | Decreased transport activity and protein expression | HeLa-mOAT1 cell | 23 |
| p.Cys427Ala | TMH10 | | | | |
| p.Cys434Ala | | | | | |

**Supplementary Table S2.** Identities and similarities scores between hOAT1 and template proteins used by automatic protein structural prediction tool I-TASSER to build initial model prior to MD simulations.

| Pairs (PDB ID) | Local sequence alignment* | | Global sequence alignment* | |
|---|---|---|---|---|
| | Identity % | Similarity % | Identity % | Similarity % |
| ***h*OAT1- *h*Glut3 (5C65)** | 100/454 (22%) | 159/454 (35%) | 126/637 (19%) | 195/637 (30.6%) |
| ***h*OAT1- *r*Glut5 (4YBQ)** | 108/477 (22.6%) | 188/477 (39.4%) | 118/645 (18.3%) | 207/645 (32.1%) |
| ***h*OAT1- XylE (4GBY)** | 105/488 (21.5%) | 187/488 (38.3%) | 112/642 (17.4%) | 198/642 (35.8%) |

** Sequence alignment was done using Needleman-Wunsch algorithm (global) and Smith-Waterman algorithm (local) using the https://www.ebi.ac.uk webserver.*

**Supplementary Table S3.** Total number of POPC ($N_{POPC}$), POPE ($N_{POPE}$), cholesterol ($N_{Chol}$), water ($N_{water}$) molecules and atoms ($N_{atoms}$) as well as initial PBC box sizes in MD simulations of substrate free (*apo*) *h*OAT1.

| **POPC:POPE:Chol ratio** | | | **1:0:0** | **3:1:0** | **3:0:1** | **2:1:1** |
|---|---|---|---|---|---|---|
| Lipids | $N_{POPC}$ | Upper leaflet | 187 | 147 | 159 | 110 |
| | | Lower leaflet | 187 | 147 | 159 | 110 |
| | | Total | 374 | 294 | 318 | 220 |
| | $N_{POPE}$ | Upper leaflet | - | 49 | - | 55 |
| | | Lower leaflet | - | 49 | - | 55 |
| | | Total | - | 98 | - | 110 |
| | $N_{chol}$ | Upper leaflet | - | - | 53 | 55 |
| | | Lower leaflet | - | - | 53 | 55 |
| | | Total | - | - | 106 | 110 |
| Water | $N_{water}$ | | 43493 | 43931 | 44340 | 44255 |
| Box size | x | | 119.99 | 120.54 | 122.16 | 121.76 |
| | y | | 120.04 | 120.66 | 125.17 | 125.25 |
| | z | | 140.47 | 140.47 | 140.47 | 140.47 |
| Natoms | | | 189376 | 192222 | 192263 | 192922 |

**Supplementary Table S4.** The list of MFS transporters used for PCA and tilt angle analysis.

| PDB ID | Transporter | Organism | Conformation | Method | Resolution | Reference |
|---|---|---|---|---|---|---|
| 4YB9 | GLUT5 | *Bos taurus* | IF | X-Ray diffraction | 3.20 Å | [1] |
| 4ZYR | LacY | *E.coli* | IF | X-Ray diffraction | 3.31 Å | [2] |
| 4GBY | XylE | *E.coli* | OF | X-Ray diffraction | 2.81 Å | [3] |
| 4GBZ | XylE | *E.coli* | $OF^{occ}$ | X-Ray diffraction | 2.89 Å | [3] |
| 4GC0 | XylE | *E.coli* | OF | X-Ray diffraction | 2.60 Å | [3] |
| 4JA3 | XylE | *E.coli* | $IF^{occ}$ | X-Ray diffraction | 3.80 Å | [4] |
| 4JA4 | XylE | *E.coli* | IF | X-Ray diffraction | 4.20 Å | [4] |
| 4QIQ | XylE | *E.coli* | IF | X-Ray diffraction | 3.51 Å | [5] |
| 4PYP | GLUT1 | *Homo sapiens* | IF | X-Ray diffraction | 3.17 Å | [6] |
| 5EQG | GLUT1 | *Homo sapiens* | IF | X-Ray diffraction | 2.90 Å | [7] |
| 5EQH | GLUT1 | *Homo sapiens* | IF | X-Ray diffraction | 2.99 Å | [7] |
| 5EQI | GLUT1 | *Homo sapiens* | IF | X-Ray diffraction | 3.00 Å | [7] |
| 4ZWB | GLUT3 | *Homo sapiens* | $OF^{occ}$ | X-Ray diffraction | 2.40 Å | [8] |
| 4ZWC | GLUT3 | *Homo sapiens* | OF | X-Ray diffraction | 2.60 Å | [8] |
| 5C65 | GLUT3 | *Homo sapiens* | OF | X-Ray diffraction | 2.65 Å | [8] |
| 4YBQ | GLUT5 | *Rattus norvegicus, Mus musculus* | OF | X-Ray diffraction | 3.27 Å | [1] |
| 4LDS | GlcP | *Staphylococcus epidermidis* | IF | X-Ray diffraction | 3.20 Å | [9] |
| 1PW4 | GlpT | *E.coli* | IF | X-Ray diffraction | 3.30 Å | [10] |
| 3WDO | YajR | *E.coli* | OF | X-Ray diffraction | 3.15 Å | [11] |
| 2GFP | EmrD | *E.coli* | OF | X-Ray diffraction | 3.50 Å | [12] |
| | OAT1 | *Homo sapiens* | IF | Alpha-fold 2 | | [13] |

**Supplementary Table S5.** Topological MFS core definition for the MFS dataset used in the present study as well as for *h*OAT1 model. For sake of readability, TMH definitions are split into (**A**) N- and (**B**) C-bundles.

(A)

| PDB ID/ System | TMH1 | TMH2 | TMH3 | TMH4 | TMH5 | TMH6 |
|---|---|---|---|---|---|---|
| 4YB9 | 26-43 | 72-91 | 101-118 | 126-146 | 163-182 | 193-211 |
| 4ZYR | 12-29 | 48-67 | 77-94 | 108-128 | 141-160 | 167-185 |
| 4GBY | 16-33 | 59-78 | 88-105 | 128-148 | 164-183 | 201-219 |
| 4GBZ | 16-33 | 59-78 | 88-105 | 128-148 | 164-183 | 201-219 |
| 4GC0 | 16-33 | 59-78 | 88-105 | 128-148 | 164-183 | 201-219 |
| 4JA3 | 16-33 | 58-77 | 87-104 | 127-147 | 166-185 | 201-219 |
| 4JA4 | 16-33 | 56-75 | 87-104 | 127-147 | 166-185 | 201-219 |
| 4QIQ | 16-33 | 56-75 | 86-103 | 127-147 | 166-185 | 200-218 |
| 4PYP | 18-35 | 63-82 | 95-112 | 120-140 | 157-176 | 187-205 |
| 5EQG | 18-35 | 63-82 | 95-112 | 120-140 | 157-176 | 187-205 |
| 5EQH | 18-35 | 63-82 | 95-112 | 120-140 | 157-176 | 187-205 |
| 5EQI | 18-35 | 63-82 | 95-112 | 120-140 | 157-176 | 187-205 |
| 4ZWB | 14-31 | 65-84 | 93-110 | 119-139 | 155-174 | 185-203 |
| 4ZWC | 14-31 | 65-84 | 93-110 | 119-139 | 155-174 | 185-203 |
| 5C65 | 14-31 | 65-84 | 93-110 | 119-139 | 155-174 | 185-203 |
| 4YBQ | 19-36 | 72-91 | 100-117 | 128-148 | 161-180 | 192-210 |
| 4LDS | 12-29 | 43-62 | 73-90 | 96-116 | 134-153 | 161-179 |
| 1PW4 | 33-50 | 64-83 | 94-111 | 121-141 | 160-179 | 189-207 |
| 3WDO | 15-32 | 50-69 | 79-96 | 104-124 | 137-156 | 165-183 |
| 2GFP | 14-31 | 44-63 | 74-91 | 98-118 | 134-153 | 158-176 |
| OAT1 | 20-37 | 132-151 | 163-180 | 191-211 | 222-241 | 249-267 |

(B)

| PDB ID/ System | TMH7 | TMH8 | TMH9 | TMH10 | TMH11 | TMH12 |
|---|---|---|---|---|---|---|
| 4YB9 | 280-302 | 316-335 | 344-361 | 375-393 | 413-433 | 440-458 |
| 4ZYR | 222-244 | 262-281 | 290-307 | 316-334 | 348-368 | 381-399 |
| 4GBY | 279-301 | 316-335 | 343-360 | 374-392 | 407-427 | 443-461 |
| 4GBZ | 279-301 | 316-335 | 343-360 | 374-392 | 407-427 | 443-461 |
| 4GC0 | 279-301 | 316-335 | 343-360 | 374-392 | 407-427 | 443-461 |
| 4JA3 | 279-301 | 316-335 | 344-361 | 372-390 | 408-428 | 444-462 |
| 4JA4 | 279-301 | 317-336 | 344-361 | 372-390 | 409-429 | 444-462 |
| 4QIQ | 281-303 | 315-334 | 344-361 | 372-390 | 409-429 | 444-462 |
| 4PYP | 275-297 | 308-327 | 337-354 | 367-385 | 405-425 | 432-450 |
| 5EQG | 275-297 | 308-327 | 337-354 | 367-385 | 405-425 | 432-450 |
| 5EQH | 275-297 | 308-327 | 337-354 | 367-385 | 405-425 | 432-450 |
| 5EQI | 275-297 | 308-327 | 337-354 | 367-385 | 405-425 | 432-450 |
| 4ZWB | 272-294 | 306-325 | 335-352 | 366-384 | 402-422 | 430-448 |
| 4ZWC | 272-294 | 306-325 | 335-352 | 366-384 | 402-422 | 430-448 |
| 5C65 | 272-294 | 306-325 | 335-352 | 367-385 | 402-422 | 430-448 |
| 4YBQ | 277-299 | 317-336 | 345-362 | 370-388 | 411-431 | 439-457 |
| 4LDS | 244-266 | 277-296 | 307-324 | 336-354 | 375-395 | 402-420 |
| 1PW4 | 254-276 | 292-311 | 323-340 | 350-368 | 386-406 | 417-435 |
| 3WDO | 214-236 | 250-269 | 280-297 | 306-324 | 340-360 | 368-386 |
| 2GFP | 207-229 | 239-258 | 267-284 | 291-309 | 326-346 | 358-376 |
| OAT1 | 335-357 | 370-389 | 397-414 | 425-443 | 457-477 | 486-504 |

**Supplementary Table S6.** The helicity of each transmembrane helix is displayed as the average of 3 replicas for each amino acid numbered from 1 as the first in a helix. Corresponding amino acids are listed in the last row.

| Number of amino acid | TM1 | TM2 | TM3 | TM4 | TM5 | TM6 | TM7 | TM8 | TM9 | TM10 | TM11 | TM12 |
|---|---|---|---|---|---|---|---|---|---|---|---|---|
| 1 | 0.0 | 0.0 | 0.0 | 0.0 | 0.0 | 0.0 | 0.0 | 0.0 | 0.0 | 0.0 | 0.0 | 0.0 |
| 2 | 0.961 | 0.331 | 0.949 | 0.992 | 0.153 | 0.352 | 0.199 | 0.963 | 0.996 | 0.541 | 0.0 | 0.702 |
| 3 | 0.963 | 0.594 | 0.988 | 1.0 | 0.338 | 0.36 | 0.295 | 0.992 | 0.999 | 0.569 | 0.0 | 0.749 |
| 4 | 0.963 | 0.862 | 0.999 | 1.0 | 0.461 | 0.972 | 0.629 | 0.996 | 1.0 | 0.917 | 0.996 | 0.749 |
| 5 | 0.358 | 0.994 | 1.0 | 1.0 | 0.517 | 0.973 | 0.634 | 0.998 | 1.0 | 0.965 | 0.996 | 0.947 |
| 6 | 0.0 | 0.979 | 1.0 | 1.0 | 0.996 | 0.999 | 0.989 | 0.999 | 1.0 | 0.953 | 1.0 | 0.994 |
| 7 | 0.0 | 0.98 | 1.0 | 1.0 | 0.999 | 0.998 | 0.99 | 0.926 | 1.0 | 0.593 | 1.0 | 0.997 |
| 8 | 0.247 | 0.96 | 1.0 | 1.0 | 0.999 | 0.841 | 1.0 | 0.915 | 1.0 | 0.209 | 1.0 | 0.998 |
| 9 | 0.533 | 0.954 | 1.0 | 1.0 | 1.0 | 0.937 | 1.0 | 0.836 | 1.0 | 0.028 | 1.0 | 0.996 |
| 10 | 0.997 | 0.999 | 1.0 | 0.999 | 0.999 | 0.996 | 1.0 | 0.017 | 1.0 | 0.571 | 1.0 | 0.825 |
| 11 | 1.0 | 1.0 | 1.0 | 1.0 | 0.994 | 0.996 | 1.0 | 0.947 | 1.0 | 0.85 | 1.0 | 0.844 |
| 12 | 1.0 | 0.923 | 1.0 | 1.0 | 0.989 | 1.0 | 1.0 | 0.979 | 1.0 | 0.891 | 1.0 | 0.948 |
| 13 | 1.0 | 0.866 | 1.0 | 1.0 | 0.986 | 0.999 | 1.0 | 0.994 | 0.999 | 0.937 | 1.0 | 0.946 |
| 14 | 1.0 | 0.995 | 0.999 | 1.0 | 0.956 | 0.992 | 1.0 | 0.999 | 0.999 | 0.964 | 0.999 | 0.874 |
| 15 | 1.0 | 0.996 | 1.0 | 1.0 | 0.928 | 0.999 | 1.0 | 0.997 | 0.997 | 0.939 | 0.995 | 0.393 |
| 16 | 0.999 | 0.999 | 1.0 | 1.0 | 0.929 | 0.997 | 1.0 | 0.992 | 0.938 | 0.928 | 0.98 | 0.0 |
| 17 | 1.0 | 0.998 | 1.0 | 1.0 | 0.756 | 0.997 | 0.999 | 0.958 | 0.867 | 0.855 | 0.96 | |
| 18 | 0.997 | 1.0 | 1.0 | 1.0 | 0.734 | 0.996 | 0.957 | 0.954 | 0.0 | 0.607 | 0.801 | |
| 19 | 0.974 | 1.0 | 0.959 | 1.0 | 0.526 | 0.943 | 0.984 | 0.65 | | 0.623 | 0.334 | |
| 20 | 0.952 | 1.0 | 0.824 | 1.0 | 0.0 | 0.853 | 0.971 | 0.634 | | 0.995 | 0.248 | |
| 21 | 0.0 | 1.0 | 0.0 | 1.0 | | 0.684 | 0.965 | 0.373 | | 0.986 | 0.947 | |
| 22 | | 0.999 | | 0.999 | | 0.0 | 0.864 | 0.0 | | 0.951 | 0.999 | |
| 23 | | 0.998 | | 0.998 | | | 0.373 | | | 0.778 | 1.0 | |
| 24 | | 0.993 | | 0.913 | | | 0.954 | | | 0.0 | 1.0 | |
| 25 | | 0.608 | | 0.487 | | | 0.997 | | | | 1.0 | |
| 26 | | 0.991 | | 0.418 | | | 0.989 | | | | 1.0 | |
| 27 | | | | 0.0 | | | 0.972 | | | | 0.996 | |
| 28 | | | | | | | | | | | 0.996 | |
| 29 | | | | | | | | | | | 0.0 | |
| Helices definition | 19-39 | 127-156 | 161-181 | 186-212 | 220-239 | 247-268 | 333-361 | 368-389 | 395-412 | 422-445 | 451-481 | 488-503 |

**Supplementary Table S7.** Sum of H-bond fractions for each pair of helices. The average along the simulations and standard deviation are provided.

| | TMH1 | TMH2 | TMH3 | TMH4 | TMH5 | TMH6 | TMH7 | TMH8 | TMH9 | TMH10 | TMH11 | TMH12 |
|---|---|---|---|---|---|---|---|---|---|---|---|---|
| TMH1 | | 0.993 ±0.334 | 0.0±0.0 | 1.493 ±0.101 | 0.227 ±0.067 | 1.107 ±0.242 | 0.41 ±0.537 | 0.0±0.0 | 0.0±0.0 | 0.0±0.0 | 0.604 ±0.32 | 0.0±0.0 |
| TMH2 | 0.993 ±0.334 | | 0.289 ±0.238 | 0.266 ±0.209 | 0.0±0.0 | 0.002 ±0.004 | 0.0±0.0 | 0.0±0.0 | 0.0±0.0 | 0.0±0.0 | 0.476 ±0.458 | 0.0±0.0 |
| TMH3 | 0.0±0.0 | 0.289 ±0.238 | | 2.678 ±0.241 | 0.0±0.0 | 1.679 ±0.469 | 0.0±0.0 | 0.0±0.0 | 0.0±0.0 | 0.0±0.0 | 0.013 ±0.009 | 0.0±0.0 |
| TMH4 | 1.493 ±0.101 | 0.266 ±0.209 | 2.678 ±0.241 | | 0.438 ±0.121 | 0.593 ±0.063 | 0.0±0.0 | 0.0±0.0 | 0.0±0.0 | 0.056 ±0.045 | 0.013 ±0.019 | 0.013 ±0.019 |
| TMH5 | 0.227 ±0.067 | 0.0±0.0 | 0.0±0.0 | 0.438 ±0.121 | | 0.266 ±0.274 | 0.612 ±0.429 | 2.582 ±0.574 | 0.0±0.0 | 1.154 ±0.194 | 0.004 ±0.003 | 0.0±0.0 |
| TMH6 | 1.107 ±0.242 | 0.002 ±0.004 | 1.679 ±0.469 | 0.593 ±0.063 | 0.266 ±0.274 | | 0.0±0.0 | 0.0±0.0 | 0.0±0.0 | 0.0±0.0 | 0.0±0.0 | 0.0±0.0 |
| TMH7 | 0.41 ±0.537 | 0.0±0.0 | 0.0±0.0 | 0.0±0.0 | 0.612 ±0.429 | 0.0±0.0 | | 0.0±0.0 | 0.609 ±0.467 | 0.512 ±0.369 | 0.316 ±0.213 | 0.014 ±0.018 |
| TMH8 | 0.0±0.0 | 0.0±0.0 | 0.0±0.0 | 0.0±0.0 | 2.582 ±0.574 | 0.0±0.0 | 0.0±0.0 | | 0.007 ±0.01 | 0.424 ±0.255 | 0.0±0.0 | 0.0±0.0 |
| TMH9 | 0.0±0.0 | 0.0±0.0 | 0.0±0.0 | 0.0±0.0 | 0.0±0.0 | 0.0±0.0 | 0.609 ±0.467 | 0.007 ±0.01 | | 2.464 ±0.668 | 0.106 ±0.045 | 1.301 ±0.577 |
| TMH10 | 0.0±0.0 | 0.0±0.0 | 0.0±0.0 | 0.056 ±0.045 | 1.154 ±0.194 | 0.0±0.0 | 0.512 ±0.369 | 0.424 ±0.255 | 2.464 ±0.668 | | 2.222 ±0.614 | 0.944 ±0.015 |
| TMH11 | 0.604 ±0.32 | 0.476 ±0.458 | 0.013 ±0.009 | 0.013 ±0.019 | 0.004 ±0.003 | 0.0±0.0 | 0.316 ±0.213 | 0.0±0.0 | 0.106 ±0.045 | 2.222 ±0.614 | | 0.043 ±0.052 |
| TMH12 | 0.0±0.0 | 0.0±0.0 | 0.0±0.0 | 0.013 ±0.019 | 0.0±0.0 | 0.0±0.0 | 0.014 ±0.018 | 0.0±0.0 | 1.301 ±0.577 | 0.944 ±0.015 | 0.043 ±0.052 | |

**Supplementary Table S8.** Inter-helical interactions captioned within a pair of the functional helices. The values represent H-bond fraction for which values above 1 highlight possible multiple hydrogen bonds. The tables represent interactions for systems: (A) POPC:POPE:Chol (2:1:1), (B) POPC:POPE (3:1), (C) POPC:Chol (3:1), (D) POPC. H-bond fraction above 0.1 were considered.

(A) POPC:POPE:Chol (2:1:1)

| | | rep1 | rep2 | rep3 |
|---|---|---|---|---|
| A-helices | TMH1 - TMH7 | - | Ala42-Tyr354 (0.856)<br>Thr36-Tyr353 (0.277) | - |
| | TMH4 - TMH10 | - | Thr445-Gly457 (0.115) | - |
| B-helices | TMH2 - TMH11 | Gln138-Ser472 (0.367)<br>Asp157-Gln455 (0.674) | - | Arg131-Glu480 (0.245) |
| | TMH5 - TMH8 | Thr224-Asn390 (1.024)<br>Ala238-Gln371 (0.846)<br>Ala241-Gln371 (0.669) | Thr224-Asn390 (1.195)<br>Ser231-Asp378 (0.817)<br>Tyr228-Leu379 (0.271)<br>Ala238-Gln371 (0.737)<br>Ser231-Lys382 (0.122) | Tyr228-Lys382 (0.119)<br>Ser231-Asp378 (0.102)<br>Ala238-Gln371 (0.786)<br>Thr224-Asn390 (0.229)<br>Ala241-Gln371 (0.124)<br>Ser231-Lys382 (0.332)<br>Ala220-Asn390 (0.104) |
| C-helices | TMH3 - TMH6 | Arg162-Phe268 (0.519)<br>Arg162-Glu270 (0.495)<br>Arg162-Trp266 (0.248) | Arg161-Ile269 (0.4)<br>Arg162-Trp266 (0.483)<br>Arg162-Phe268 (0.432)<br>Arg162-Glu270 (0.577)<br>Arg161-Glu270 (0.334) | Arg162-Trp266 (0.669)<br>Arg162-Glu270 (0.534) |
| | TMH9 - TMH12 | Arg395-Leu504 (0.397)<br>Arg395-Val502 (0.151 | Arg395-Leu504 (0.56)<br>Gln398-Thr501 (0.302)<br>Arg395-Thr501 (0.385) | Arg395-Leu503 (1.12)<br>Arg394-Leu504 (0.845) |

(B) POPC:POPE (3:1)

| | | rep1 | rep2 | rep3 |
|---|---|---|---|---|
| A-helices | TMH1 - TMH7 | - | Asn39-Tyr353 (0.227)<br>Asn39-Tyr354 (0.576) | - |
| | TMH4 - TMH10 | - | - | - |
| B-helices | TMH2 - TMH11 | Asp157-Gln455 (0.185) | Gln135-Pro473 (0.213)<br>Gln138-Ser472 (0.157)<br>Phe152-Gln455 (0.516) | Gln138-Ser476 (0.626)<br>Asp157-Gln455 (0.214) |
| | TMH5 - TMH8 | Ser231-Asp378 (0.895)<br>Leu237-Gln371 (0.795)<br>Gly227-Lys382 (0.249) | Gln234-Phe374 (0.222)<br>Ser231-Asp378 (0.197)<br>Ala238-Gln371 (0.287)<br>Gly227-Lys382 (0.203)<br>Ala241-Gln371 (0.104) | Ser231-Asp378 (0.966)<br>Tyr242-Gln371 (0.333)<br>Gln234-Asp378 (0.241) |
| C-helices | TMH3 - TMH6 | Arg162-Trp266 (0.704)<br>Arg162-Phe268 (0.297)<br>Arg162-Glu270 (0.398)<br>Arg161-Glu270 (0.252) | Arg162-Trp266 (0.654)<br>Arg162-Glu270 (0.766)<br>Arg162-Phe268 (0.432)<br>Tyr169-Phe258 (0.218)<br>Tyr169-Phe259 (0.12) | Arg162-Trp266 (0.728)<br>Arg162-Glu270 (0.674)<br>Arg162-Phe268 (0.306)<br>Lys163-Glu270 (0.14) |
| | TMH9 - TMH12 | Arg395-Leu504 (0.774)<br>Gln398-Thr501 (0.344)<br>Met399-Ser498 (0.221) | Arg395-Leu504 (0.827)<br>Gln398-Thr501 (0.218)<br>Met399-Ser498 (0.134) | Arg395-Leu504 (0.603)<br>Arg395-Thr501 (0.114) |

(C) POPC:Chol (3:1)

| | | rep1 | rep2 | rep3 |
|---|---|---|---|---|
| A-helices | TMH1 - TMH7 | - | Asn39-Tyr353 (0.261)<br>Asn39-Tyr354 (0.17)<br>Asn35-Tyr354 (0.439) | Asn39-Tyr354 (0.261) |
| | TMH4 - TMH10 | - | Hie217-Glu447 (0.419) | - |
| B-helices | TMH2 - TMH11 | Gly153-Thr456 (0.56)<br>Asp157-Gln455 (0.116) | Arg134-Glu480 (1.196)<br>Gln138-Ser476 (0.304)<br>Gln135-Glu480 (0.323)<br>Gln135-Ser476 (0.132)<br>Arg131-Glu480 (0.127) | Gln135-Pro473 (0.185)<br>Asp157-Gln455 (0.414) |
| | TMH5 - TMH8 | Thr224-Asn390 (1.068)<br>Ser231-Asp378 (0.941)<br>Ala238-Gln371 (0.425) | Ser231-Asp378 (0.966)<br>Thr224-Asn390 (0.943)<br>Tyr228-Leu379 (0.262)<br>Ala238-Gln371 (0.717) | Ser231-Asp378 (0.967)<br>Tyr228-Lys382 (0.452)<br>Leu237-Gln371 (0.71) |
| C-helices | TMH3 - TMH6 | Tyr169-Phe259 (0.212)<br>Arg162-Glu270 (0.693)<br>Arg162-Trp266 (0.205) | Tyr169-Phe262 (0.591)<br>Arg162-Trp266 (0.435)<br>Arg162-Glu270 (0.825)<br>Tyr169-Ser265 (0.221)<br>Arg162-Phe268 (0.239) | Arg162-Phe268 (0.771)<br>Arg162-Glu270 (0.764)<br>Lys163-Glu270 (0.37) |
| | TMH9 - TMH12 | Arg395-Leu504 (0.728)<br>Arg395-Val502 (0.158) | Arg395-Leu504 (1.347) | Arg395-Leu504 (0.92) |

(D) POPC

| | | rep1 | rep2 | rep3 |
|---|---|---|---|---|
| A-helices | TMH1 - TMH7 | - | - | Asn35-Tyr354 (0.251) |
| | TMH4 - TMH10 | - | - | - |
| B-helices | TMH2 - TMH11 | Gln138-Pro473 (0.664)<br>Arg134-Glu480 (1.366)<br>Gln135-Glu480 (0.674)<br>Gln135-Ser476 (0.3) | Asp157-Gln455 (0.627) | Val145-Arg466 (0.525)<br>Gly153-Thr456 (0.158)<br>Ala156-Gln455 (0.157) |
| | TMH5 - TMH8 | Ser231-Asp378 (0.966)<br>Ala238-Gln371 (0.619) | Tyr228-Leu379 (0.678)<br>Ala238-Gln371 (0.208)<br>Gly239-Gln371 (0.148)<br>Thr224-Lys382 (0.231) | Thr224-Asn390 (0.954)<br>Tyr228-Leu379 (0.383)<br>Ser231-Asp378 (0.545)<br>Ala238-Gln371 (0.397)<br>Ser231-Lys382 (0.128) |
| C-helices | TMH3 - TMH6 | Arg162-Trp266 (0.465)<br>Arg162-Phe268 (0.376)<br>Arg162-Glu270 (0.349)<br>Arg161-Glu270 (0.47)<br>Arg162-Phe267 (0.126) | Arg162-Glu270 (0.561)<br>Arg162-Trp266 (0.465)<br>Arg162-Phe268 (0.108) | Arg162-Phe268 (0.613)<br>Arg162-Glu270 (0.757)<br>Arg162-Trp266 (0.292) |
| | TMH9 - TMH12 | Arg395-Leu504 (0.785) | Arg395-Leu504 (0.77) | Arg395-Leu504 (1.076)<br>Gln417-Met485 (0.144) |

**Supplementary Table S9.** The hydrogen bonding network of charge-relay system split into interactions within N-bundle triad, C-bundle triad, between bundles (inter-bundle/inter-triad) and interaction of motifs with TMH11, TMH5 and TMH4. **(A)** The interactions averaged over 3 replicas of *h*OAT1 in OF state **(B)** interactions averaged over 3 replicas of *h*OAT1 (AF2) in IF state. Values above 1.0 represent possible multiple hydrogen bonds.

(A)

| | | **N-bundle** | |
|---|---|---|---|
| | A-motif | E[X6]R | [P/X]ESRW[L/X] |
| A-motif | - | **Arg161-Glu212** (1.518) | Arg162-Glu270 (0.569)<br>Arg161-Ile269 (0.126)<br>**Arg161-Glu270** (0.109) |
| E[X6]R | - | - | Glu212-Arg273 (1.16)<br>**Glu212-Ser271** (0.620) |
| [P/X]ESRW[L/X] | - | - | - |
| | | **C-bundle** | |
| | A-motif | E[X6]R | [P/X]ESRW[L/X] |
| A-motif | - | **Arg394-Glu447** (1.129) | **Arg394-Glu506** (0.674)<br>**Arg395-Glu506** (0.322) |
| E[X6]R | - | - | Glu447-Thr507 (0.630) |
| [P/X]ESRW[L/X] | - | - | - |
| | | **Inter-bundle** | |
| | A-motif | E[X6]R | [P/X]ESRW[L/X] |
| A-motif | - | - | - |
| E[X6]R | - | Arg219-Glu447 (1.334) | - |
| [P/X]ESRW[L/X] | - | - | - |
| | | **Motifs to TMH11/5/4** | |
| | TMH11 | TMH5 | TMH4 |
| A-motif | Asp157-Gln455 (0.224) | Thr224-Asn390 (0.776) | Arg161-Asn205 (0.207) |
| E[X6]R | Glu212-Gln455 (0.464) | Ala220-Asn390 (0.134) | - |

**(B)**

| | | | |
|---|---|---|---|
| | **N-bundle** | | |
| | A-motif | E[X6]R | [P/X]ESRW[L/X] |
| A-motif | - | Arg161-Glu212 (1.984) | Arg162-Glu270 (0.738)<br>Arg161-Glu270 (0.351)<br>Asp157-Trp274 (0.707) |
| E[X6]R | - | - | Trp213-Glu270 (0.186)<br>Trp213-Ile269 (0.126)<br>Glu212-Ser271 (0.997)<br>Glu212-Arg273 (0.194) |
| [P/X]ESRW[L/X] | - | - | - |
| | **C-bundle** | | |
| | A-motif | E[X6]R | [P/X]ESRW[L/X] |
| A-motif | - | Arg394-Glu447 (1.948) | Arg394-Glu506 (0.821)<br>Arg395-Glu506 (0.532) |
| E[X6]R | - | - | Glu447-Thr507 (0.982) |
| [P/X]ESRW[L/X] | - | - | - |
| | **Inter-bundle** | | |
| | A-motif | E[X6]R | [P/X]ESRW[L/X] |
| A-motif | - | - | - |
| E[X6]R | - | - | - |
| [P/X]ESRW[L/X] | - | - | - |
| | **Motifs to TMH11/5/4** | | |
| | TMH11 | TMH5 | TMH4 |
| A-motif | - | - | - |
| E[X6]R | - | - | - |

#### 2. Supplemental Figures

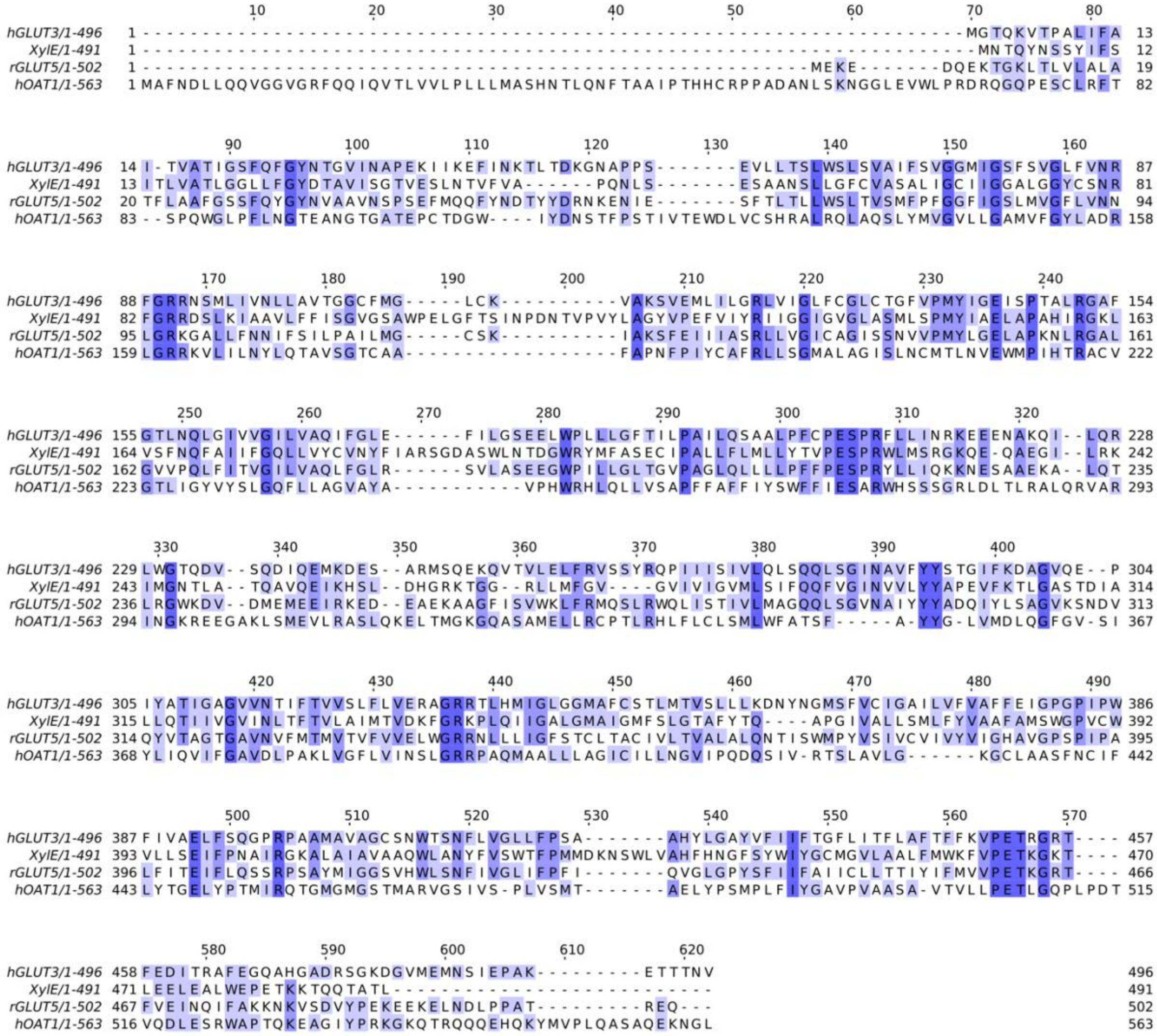

**Supplementary Figure S1.** Sequence alignment of *h*OAT1, *h*Glut3 (PDBID: 5C65), *r*Glut5 (PDBID: 4YBQ), XylE (PDBID: 4GBY) using clustal omega algorithm. Picture was rendered using JalView 2.1 with conserved amino acid colored in a blue gradient from similar (light) to identical (dark).

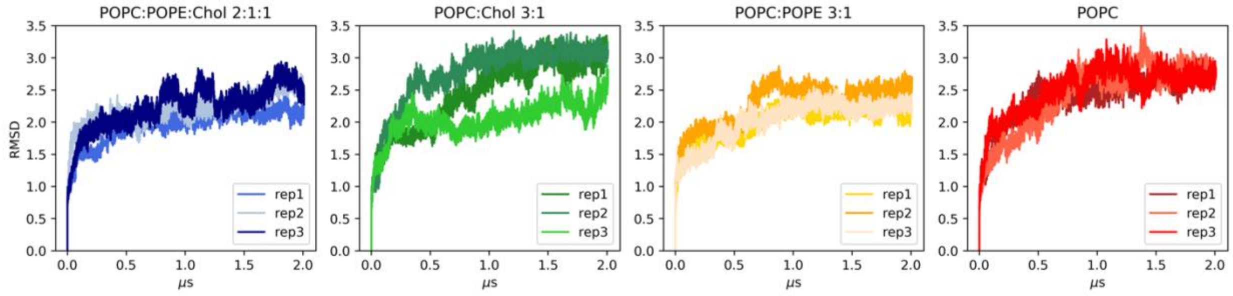

**Supplementary Figure S2.** The plot of root means square deviation (Å) in POPC:POPE:Chol (2:1:1), POPC:Chol (3:1), POPC:POPE (3:1) and pure POPC membrane in time evolution (μs).

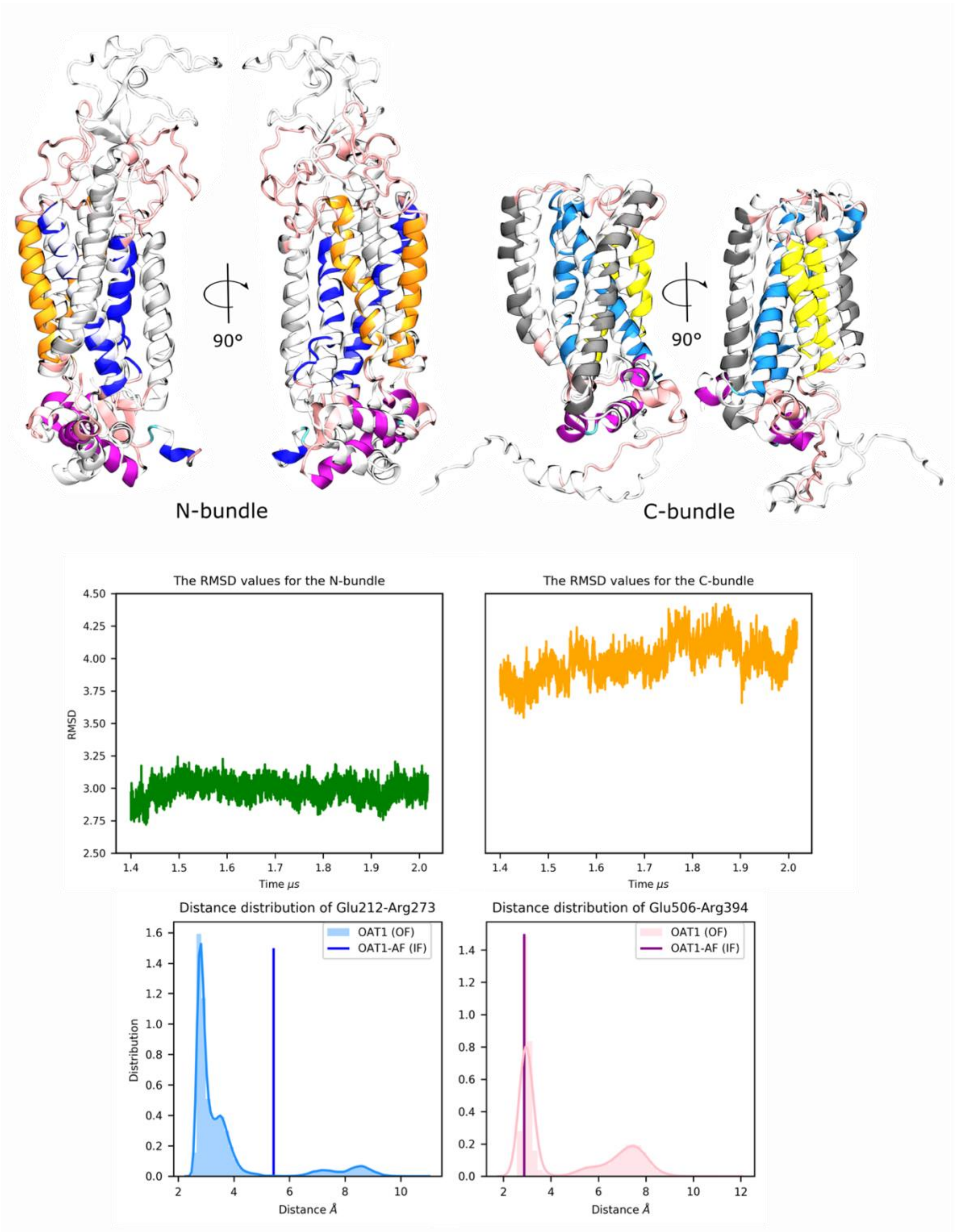

**Supplementary Figure S3.** The comparison of AF2 (IF) and present study *h*OAT1 (OF) models. The alignment is displayed by a bundle, similarly like RMSD plots below. The crucial distributions for conformation distances are plotted for present *h*OAT1 model with vertical line representing distance for AF2 model.

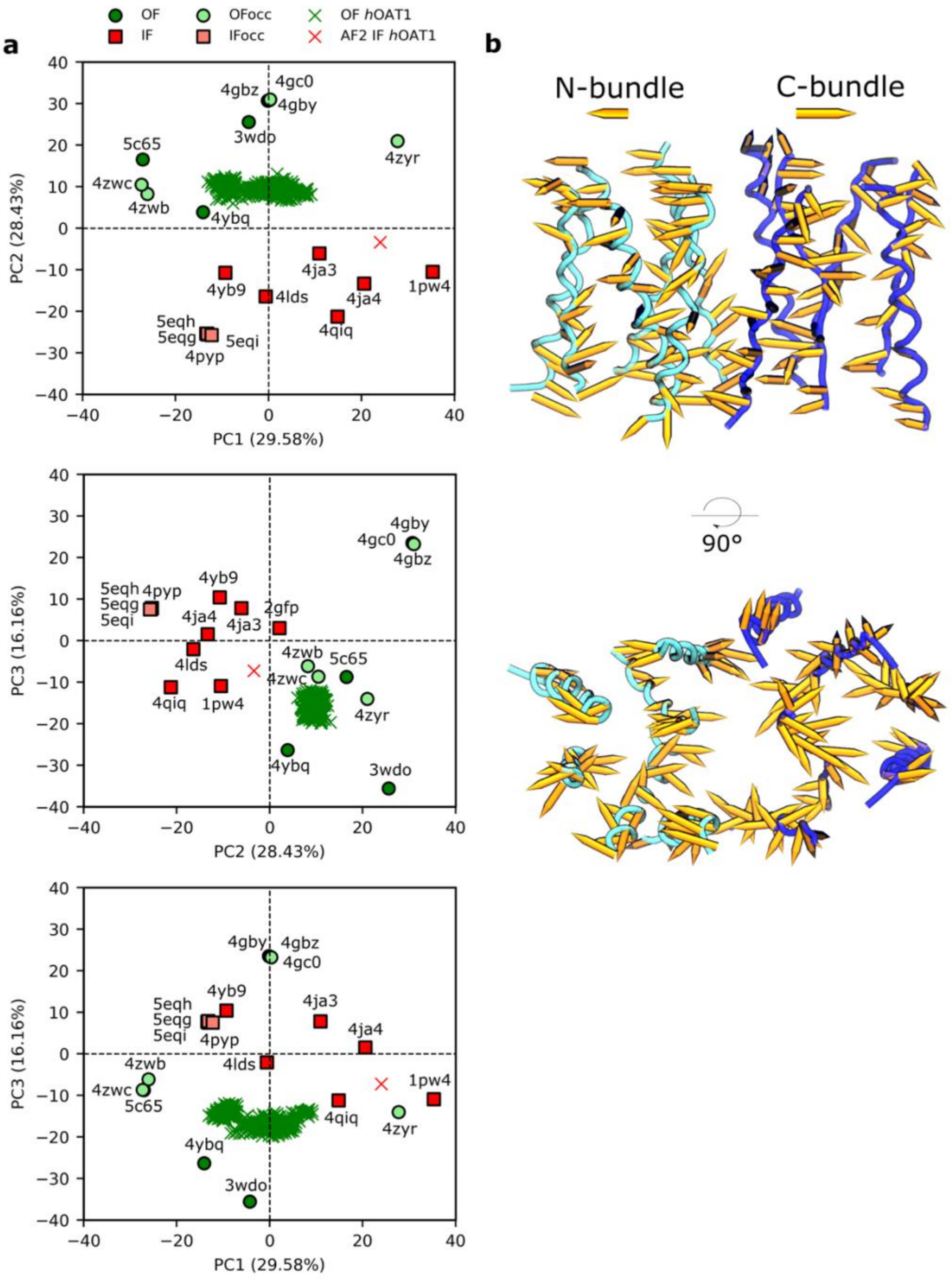

**Supplementary Figure S4.** (a) *h*OAT1 (both conformations) projected onto MFS conformational space formed by resolved MFS proteins. (b) Porcupine plot presenting PC2 (distinguishes OF and IF conformation) shows higher variability for C-bundle.

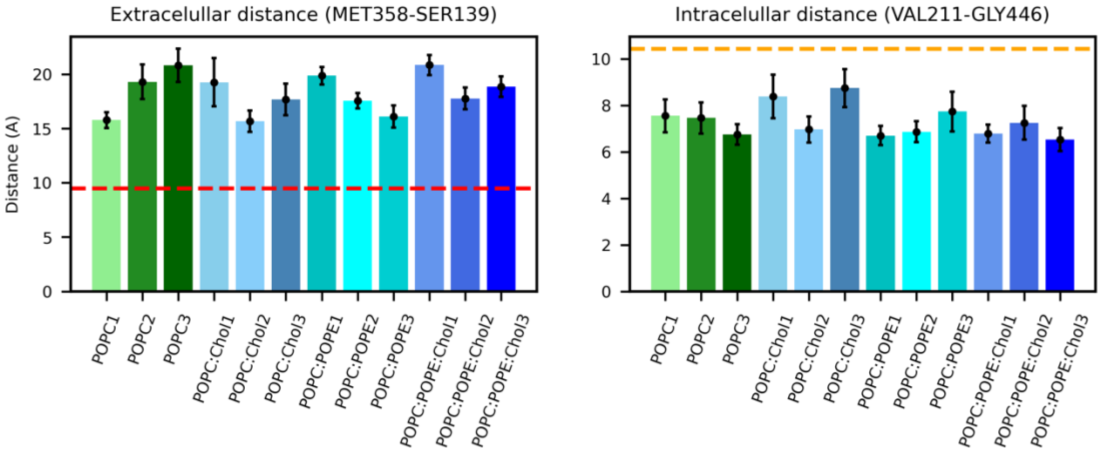

**Supplementary Figure S5.** Barplot representing distances of extracellular (Met358-Ser139) and intracellular (Val211-Gly446) distances stressed out and compared with previous *h*OAT1 model in IF conformation by Tsigelny *et al.* [24].

(A) POPC:POPE:Chol 2:1:1

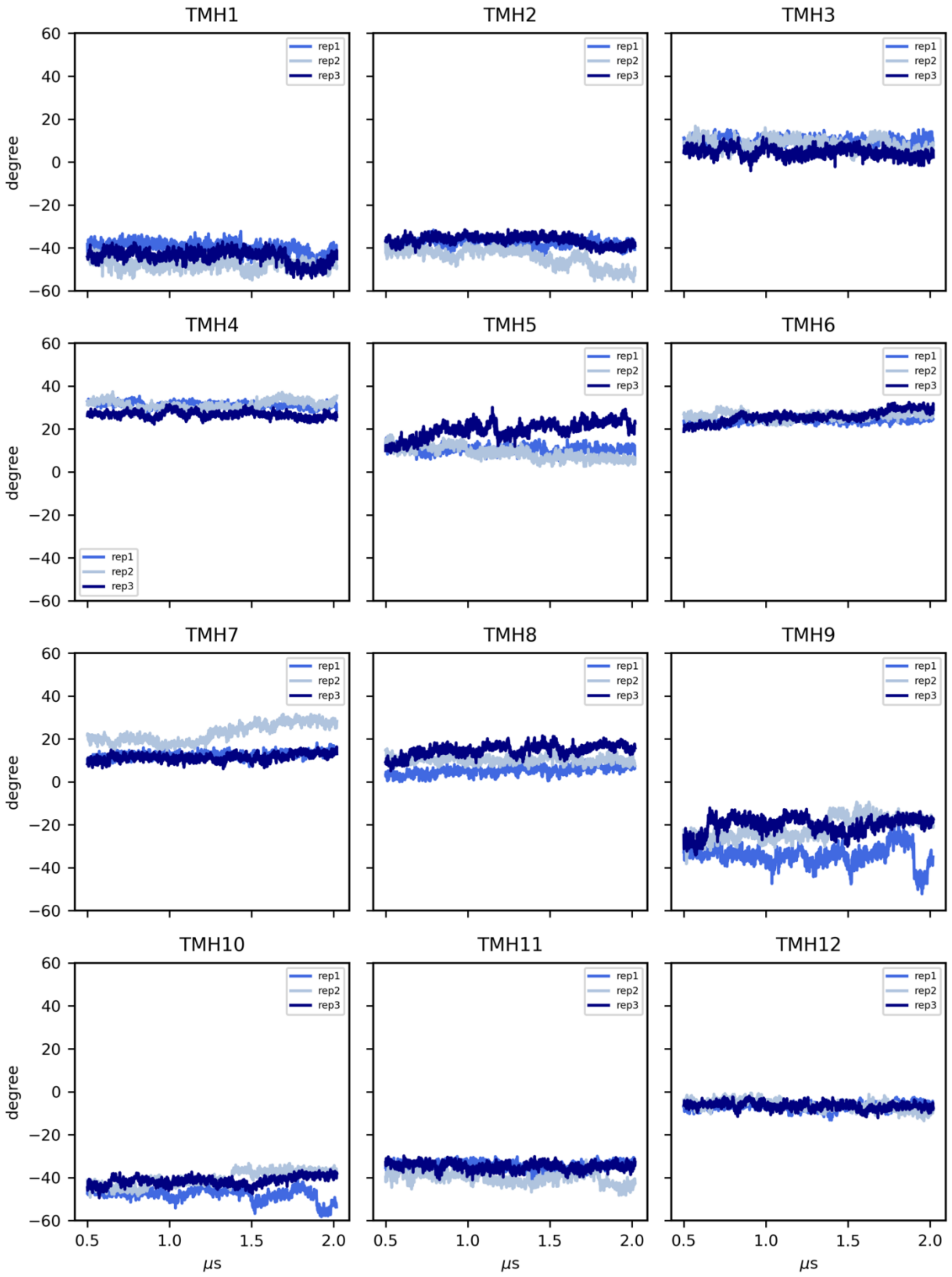

(B) POPC:Chol 3:1

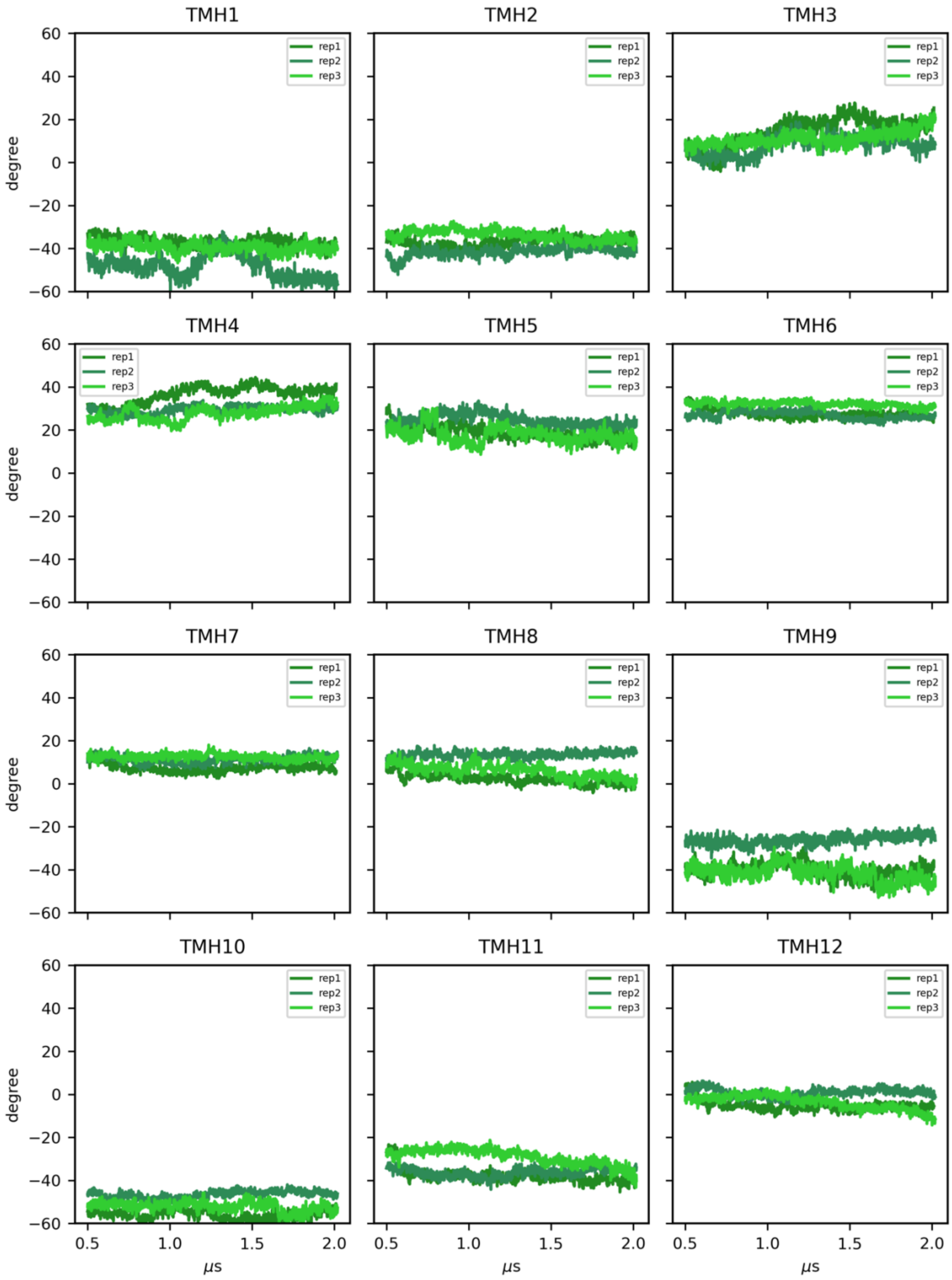

(C) POPC:POPE 3:1

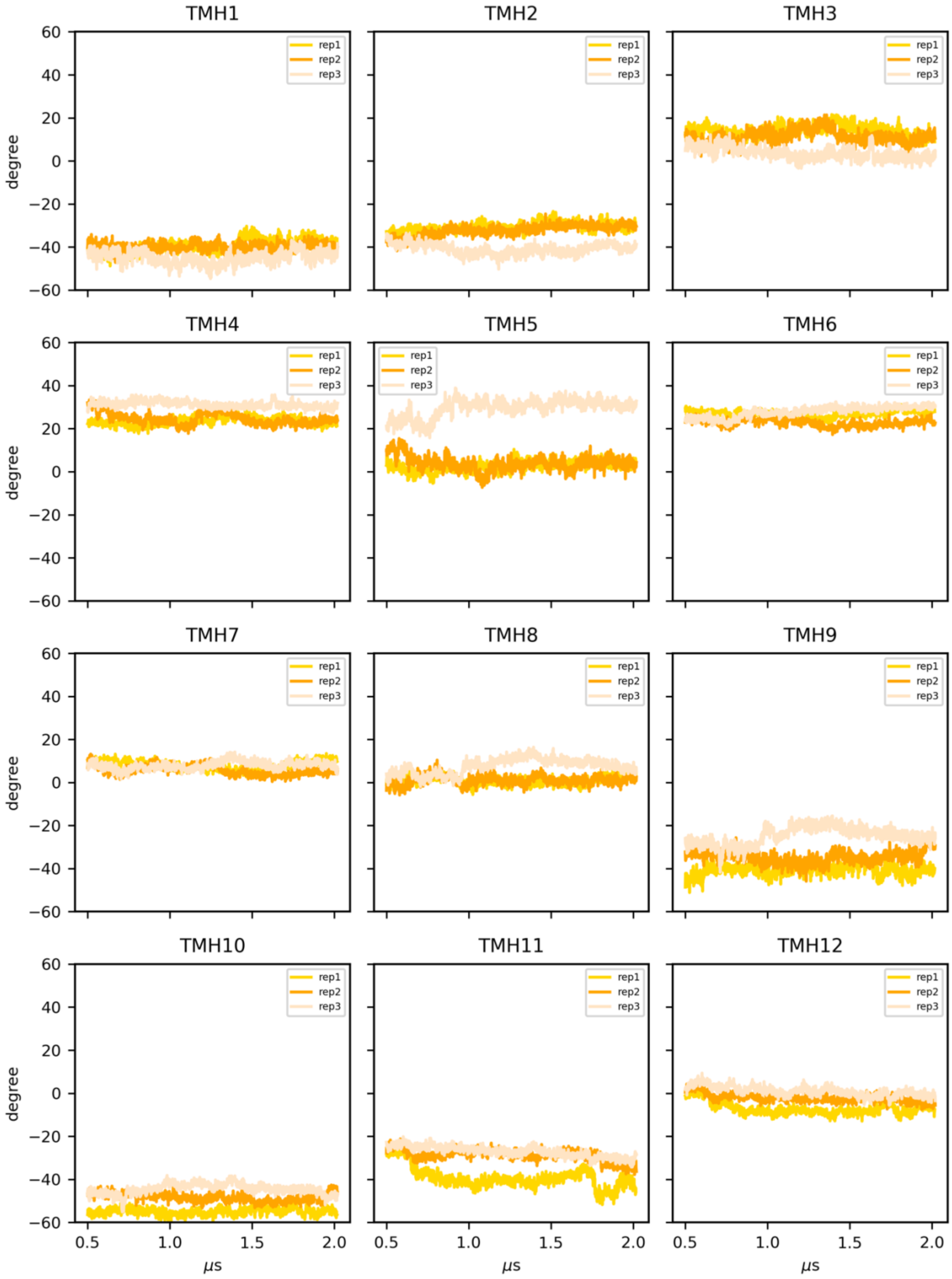

(D) POPC

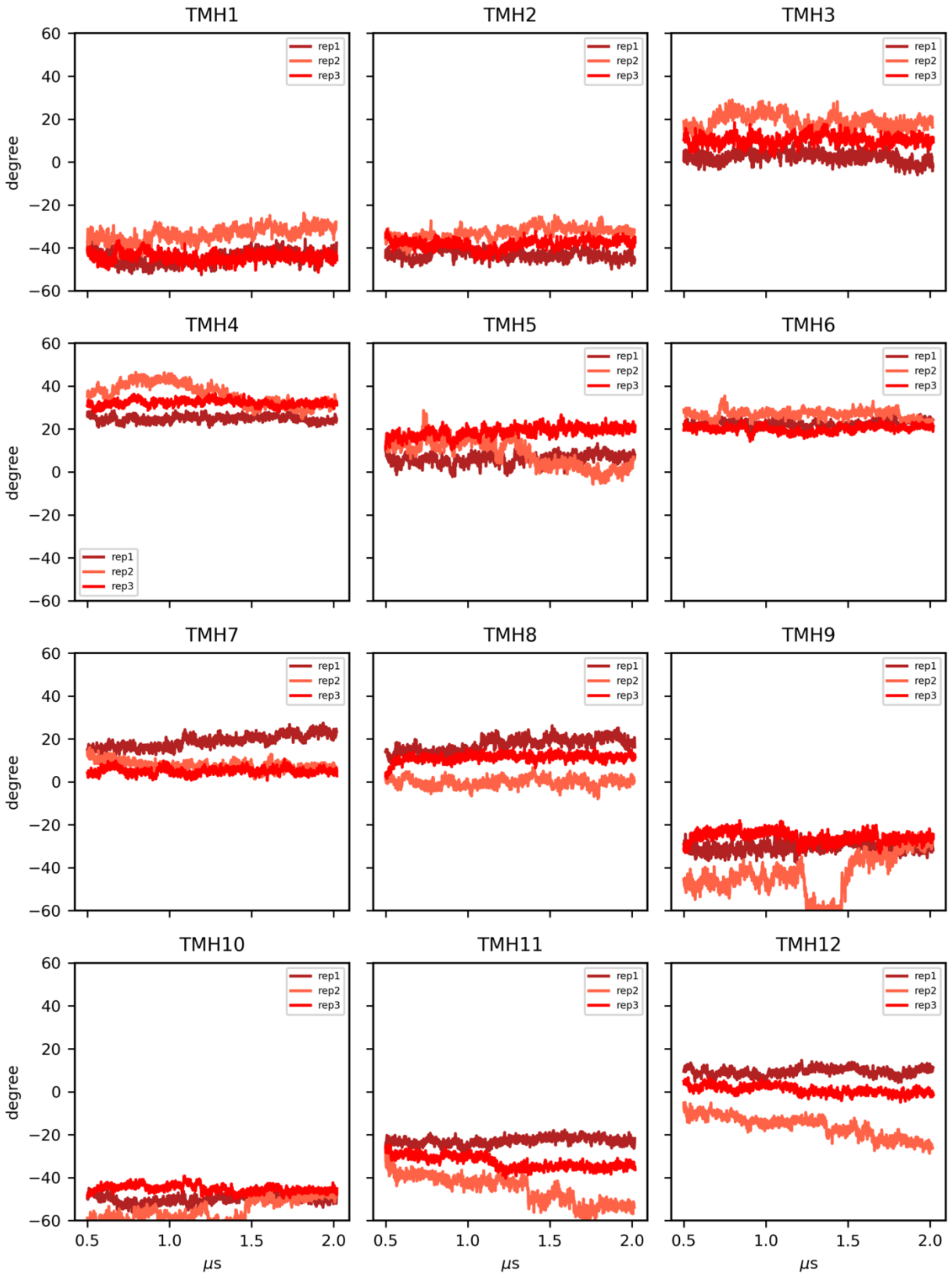
TMH1
TMH2
TMH3
TMH4
TMH5
TMH6
TMH7
TMH8
TMH9
TMH10
TMH11
TMH12
rep1
rep2
rep3
degree
μs
60
40
20
0
−20
−40
−60
0.5
1.0
1.5
2.0

**Supplementary Figure S6.** The evolution of tilt angles (clock-wise) divided by transmembrane helices of equilibrated trajectory in time for (A) POPC:POPE:Chol (2:1:1), (B) POPC:Chol (3:1), (C) POPC:POPE (3:1), (D) POPC.

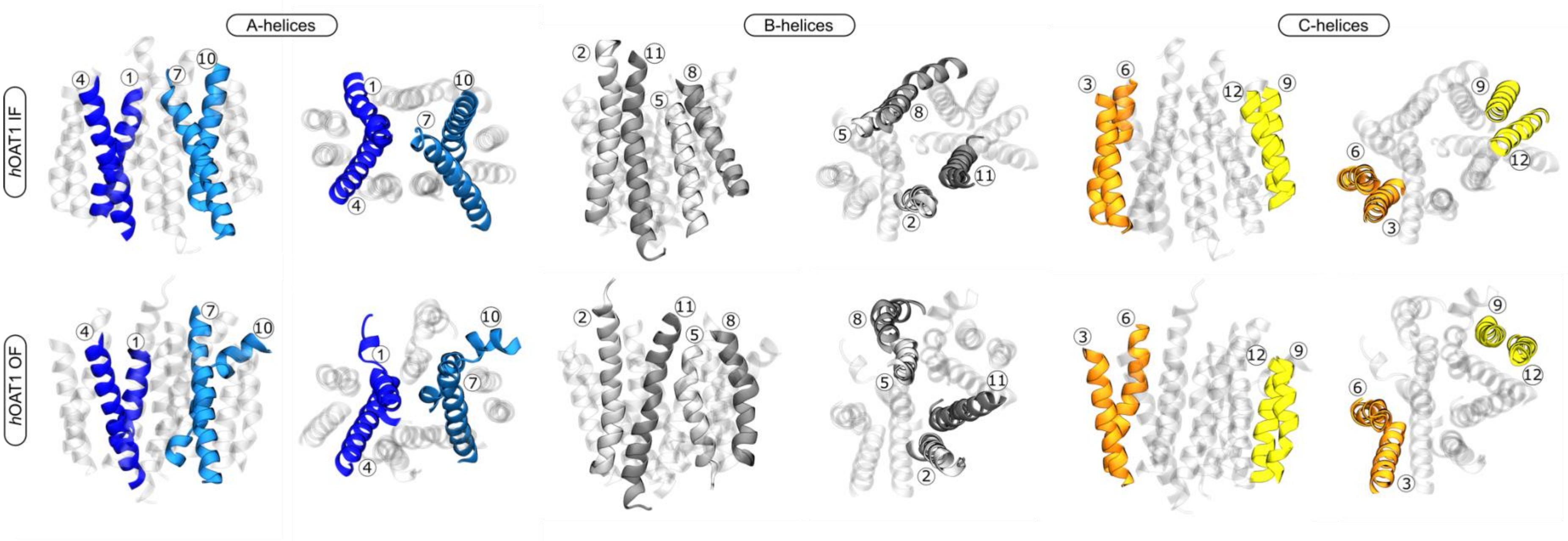

**Supplementary Figure S7.** Comparison of A-, B-, and C-helices in AF2 (upper) and present study model (bottom) of *h*OAT1.

(A) POPC:POPE:Chol 2:1:1 *h*OAT1 OF

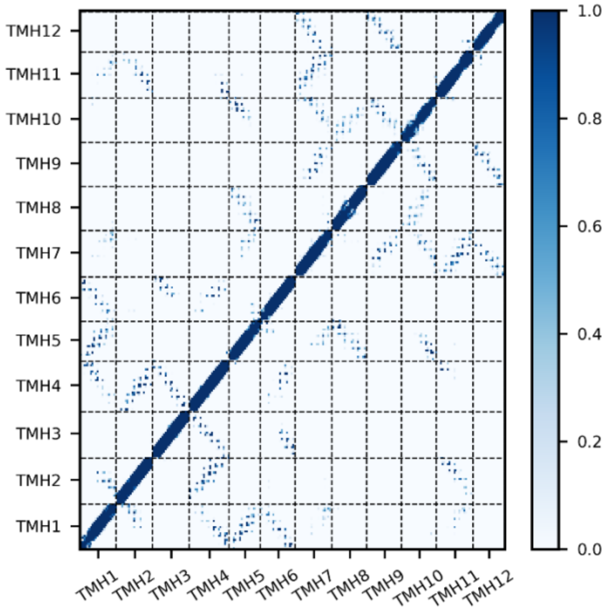

(B) POPC:POPE:Chol 2:1:1 *h*OAT1 IF

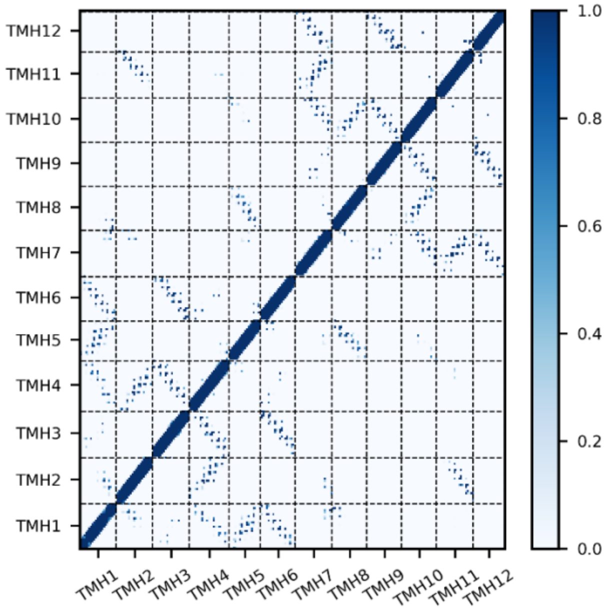

**Supplementary Figure S8.** The averaged along MD simulations contact maps of *h*OAT1 in (A) OF and (B) IF conformation divided by transmembrane helices.

(A) *h*OAT1 OF in POPC:POPE:Chol

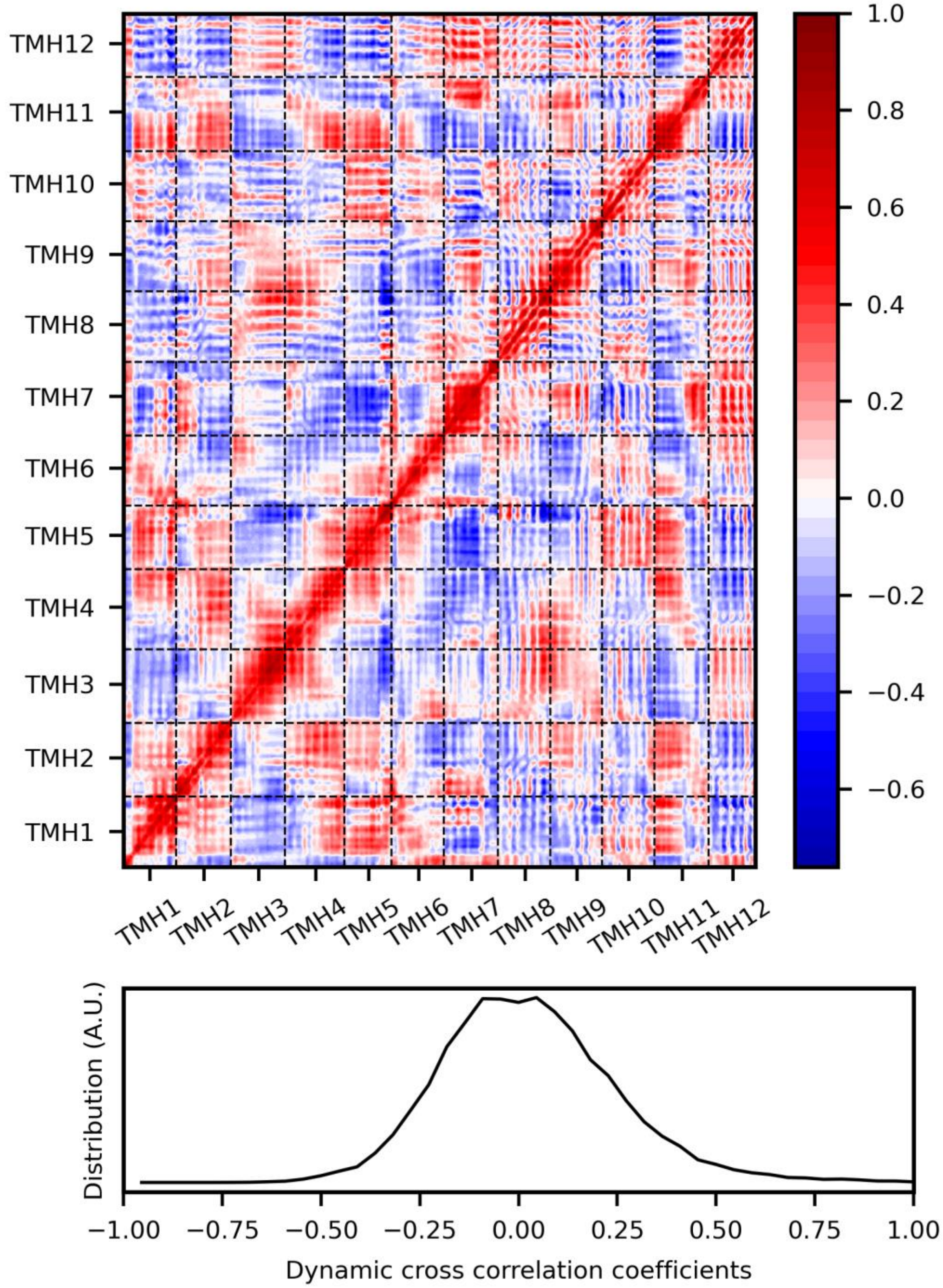

(B) *h*OAT1 IF in POPC:POPE:Chol (2:1:1)

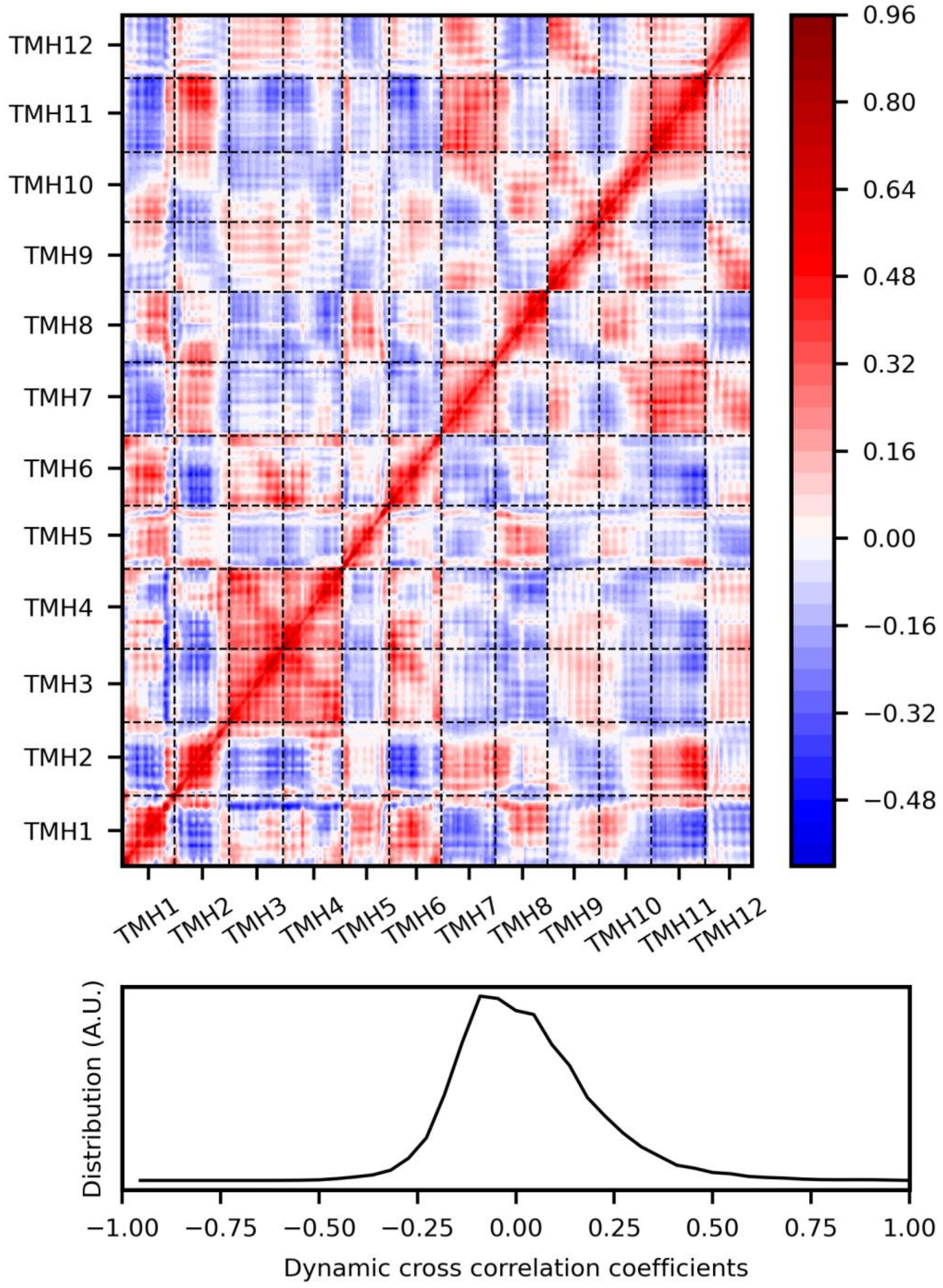

**Supplementary Figure S9.** Dynamic Cross Correlation Matrix (DCCM) of (A) *h*OAT1 OF and (B) *h*OAT1 IF (in POPC:POPE:Chol 2:1:1) divided by transmembrane helices.

(A) POPC:POPE:Chol (2:1:1)

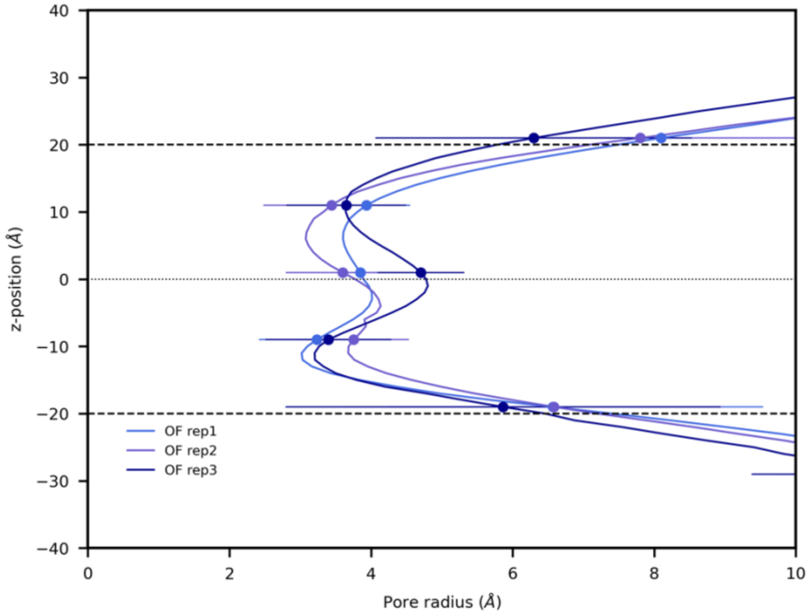

(B) POPC:Chol (3:1)

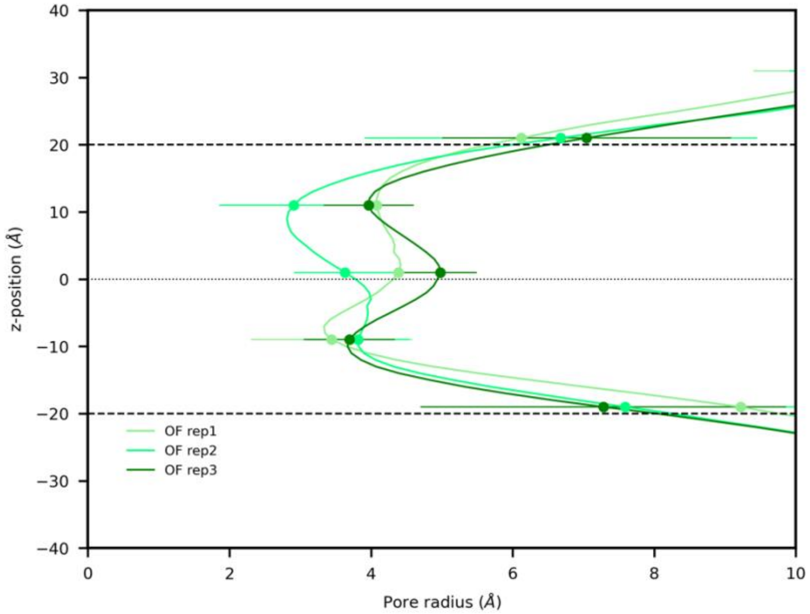

(C) POPC:POPE (3:1)

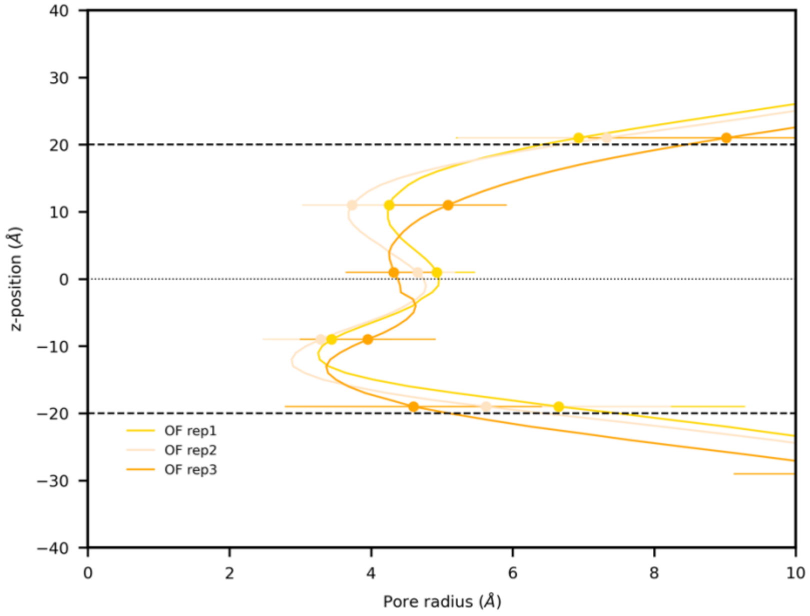

(D) POPC

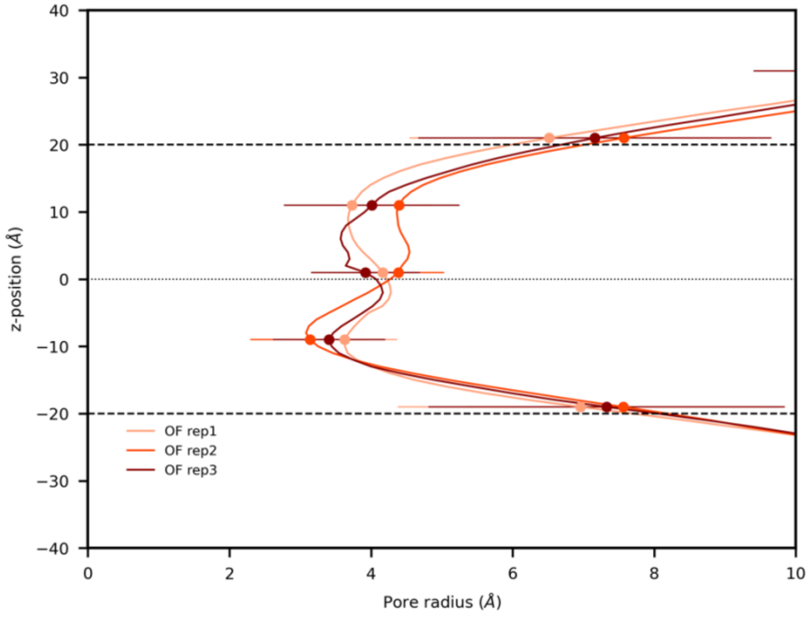

**Supplementary Figure S10.** Cavity pore radii of *h*OAT1 averaged over 500 snapshots of equilibrated trajectory over z axis for (A) POPC:POPE:Chol (2:1:1); (B) POPC:Chol (3:1); (C) POPC:POPE (3:1); (D) POPC.

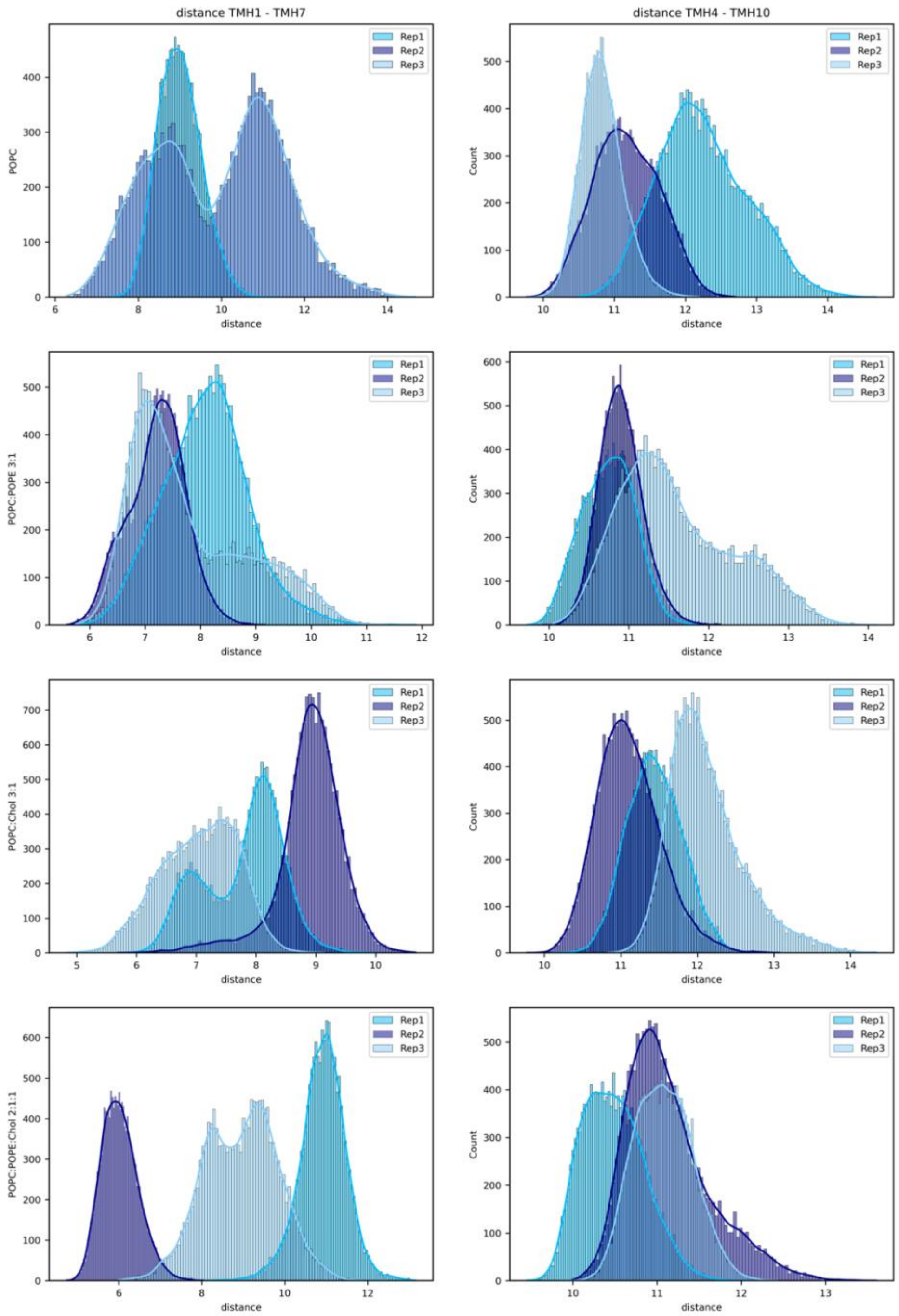

**Supplementary Figure S11.** Distribution of intra- (TM4-TM10; right column) and extracellular (TM7-TM10; left column) distances. The distances were considered as centers of mass (only backbones) of following residues: 36-40, 354-357 for TMH1-TMH7 and 207-210; 442-445 for TMH4-TMH10.

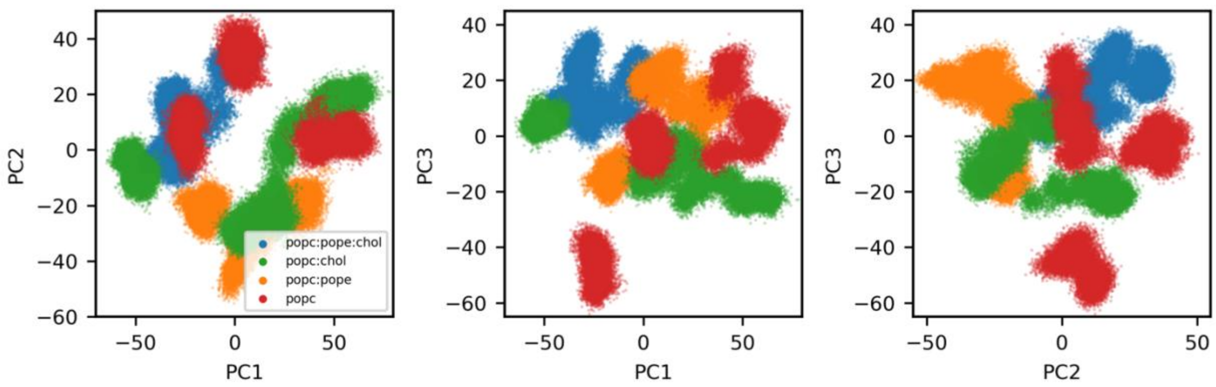

**Supplementary Figure S12.** Conformational space obtained by PCA. Projected PC1 and PC2 represent 39% and 19% of variability.

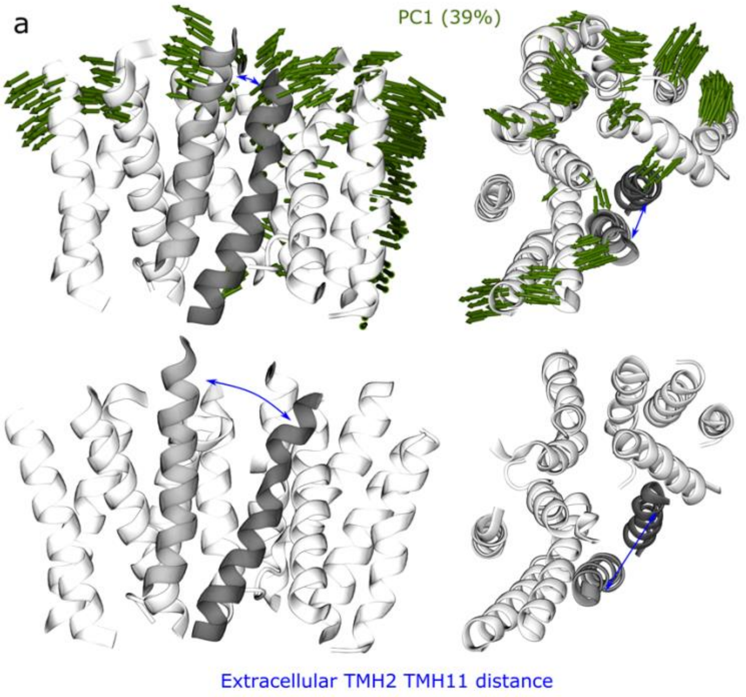

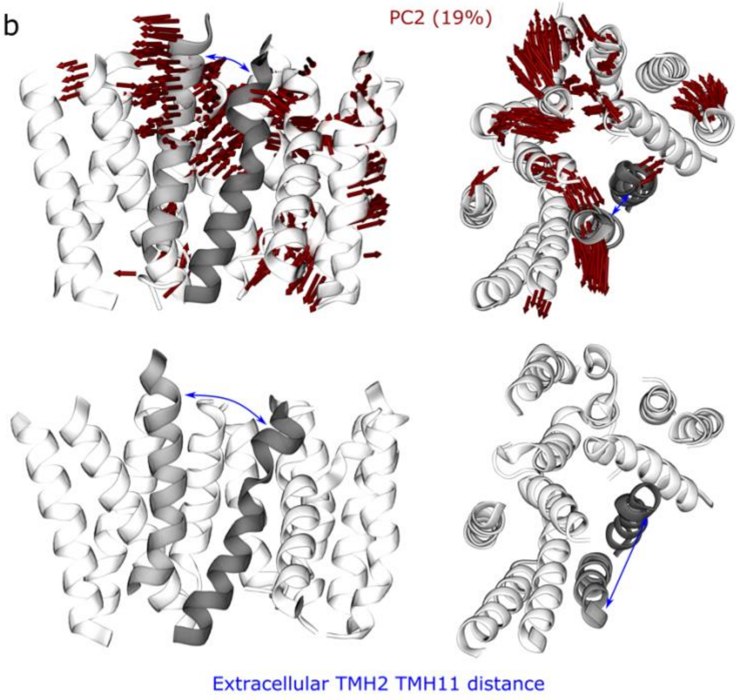

**Supplementary Figure S13.** Visualization of (a) PC1 and (b) PC2 (from Supplementary Figure S12) representing opening of the extracellular cavity of *h*OAT1. Both PCs display correlated movement of TMH2 (light grey) and TMH11 (dark grey).

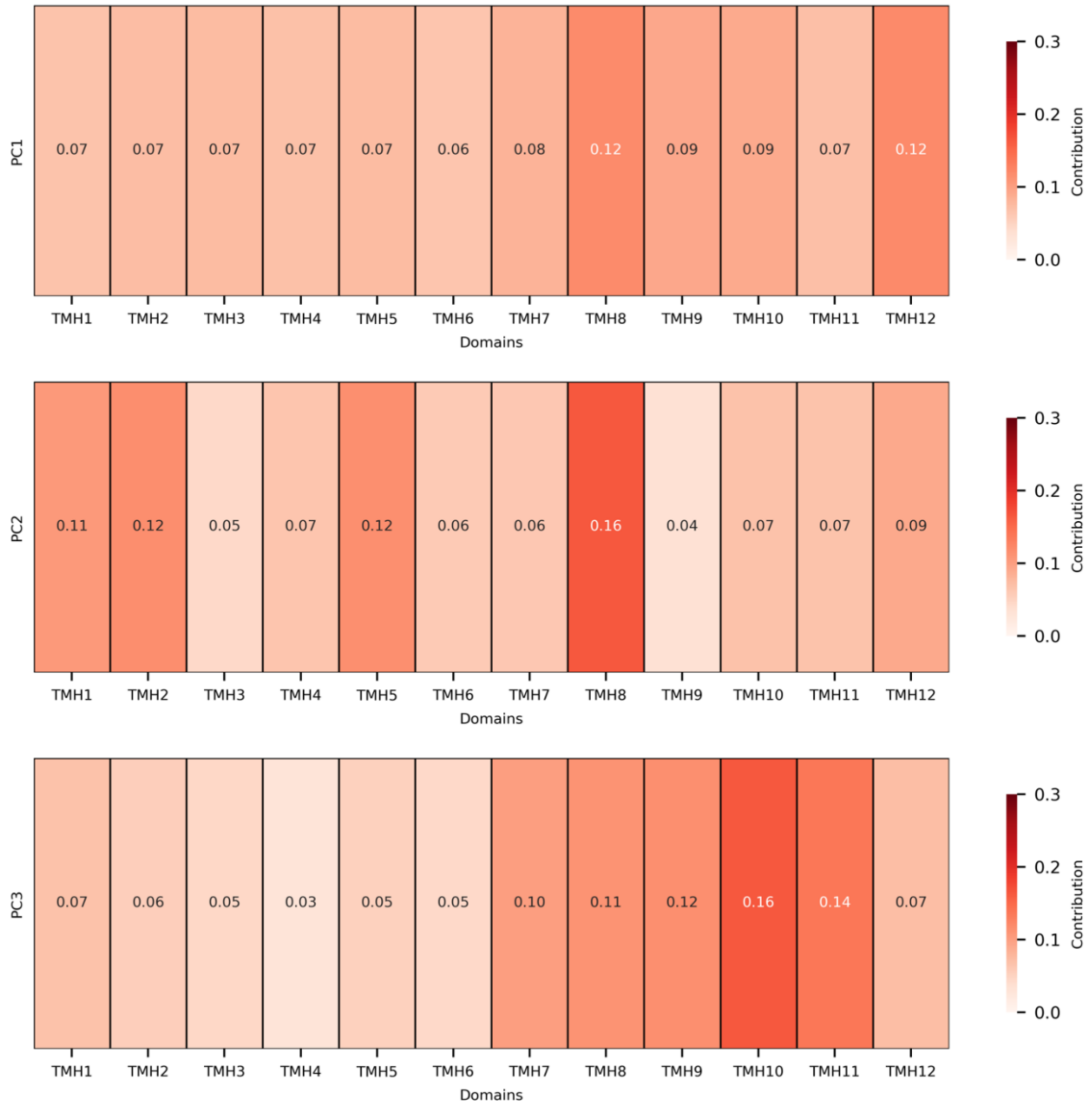

**Supplementary Figure S14.** The contribution of each TMH into PC1, PC2 and PC3.

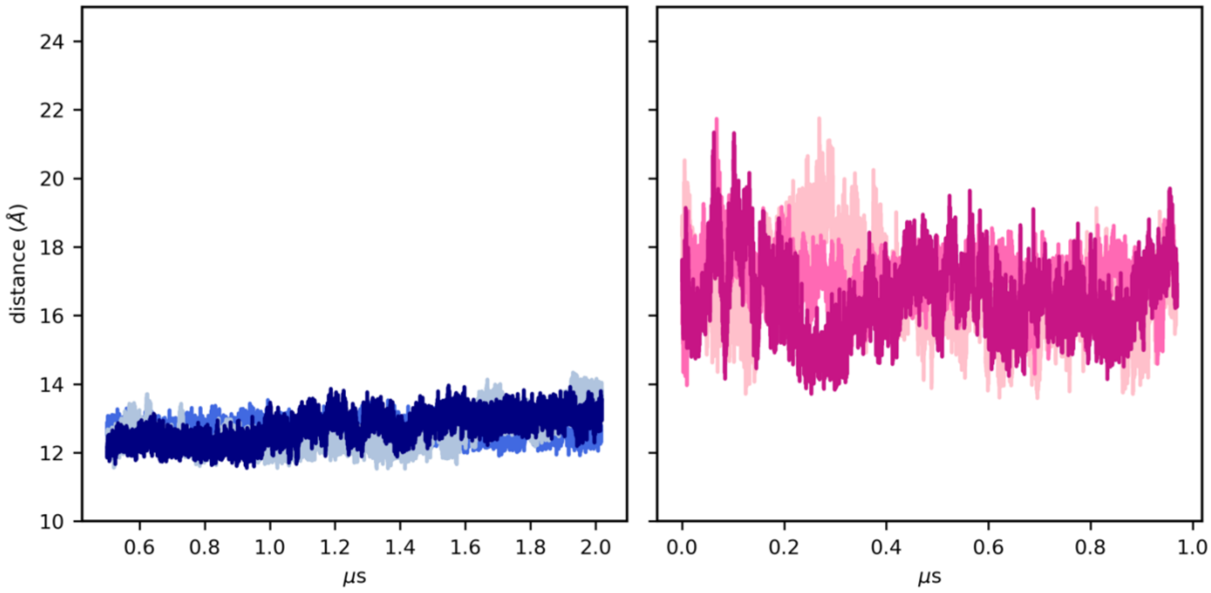

**Supplementary Figure S15.** Inter-bundle distance of E[$X_6$]R motifs in N- and C-bundle pictured along MD simulations of OF (left) and IF (right) *h*OAT1.

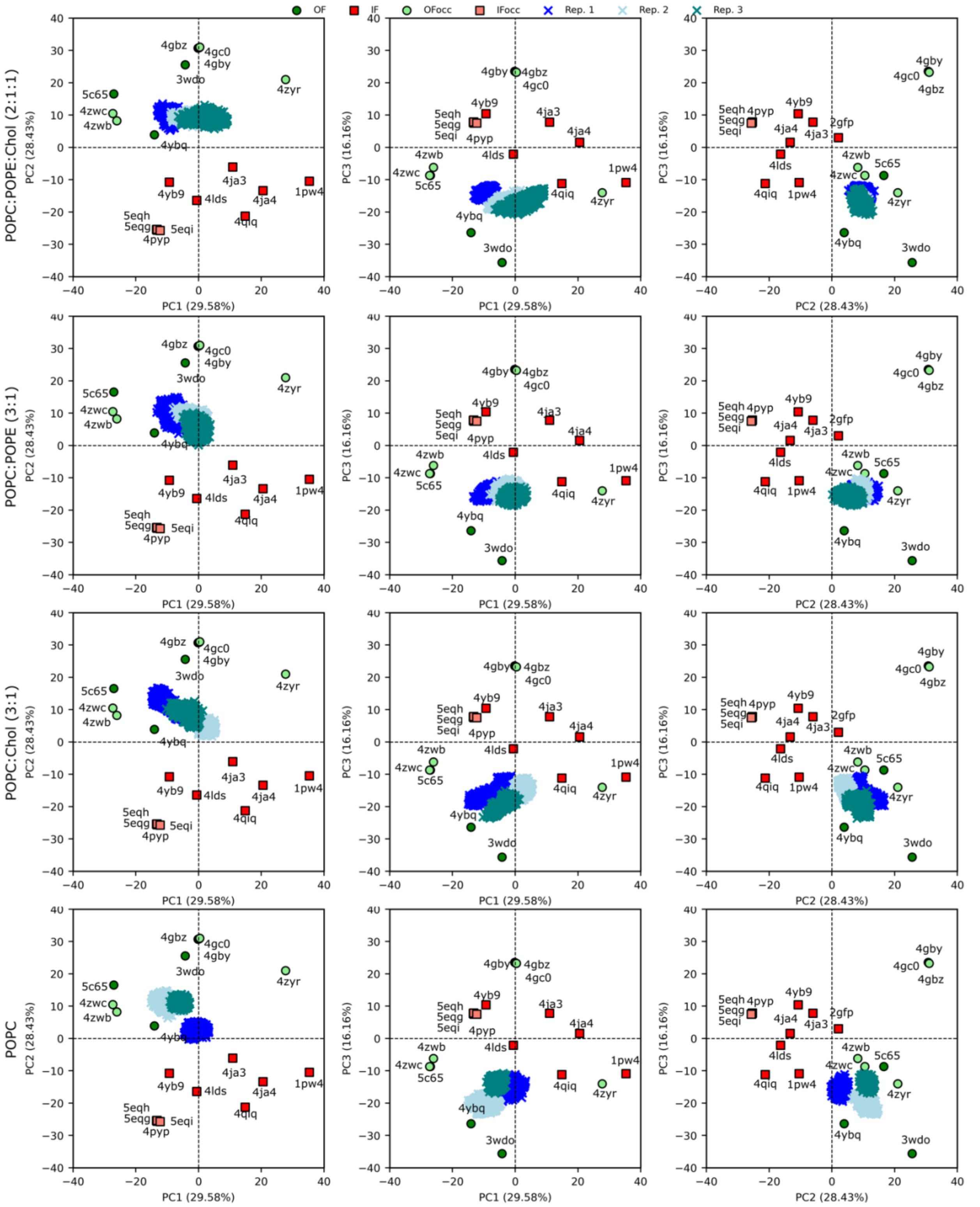

**Supplementary Figure S16.** The PCA projection of MFS conformational space including *h*OAT1 (OF) models in different lipid bilayers.

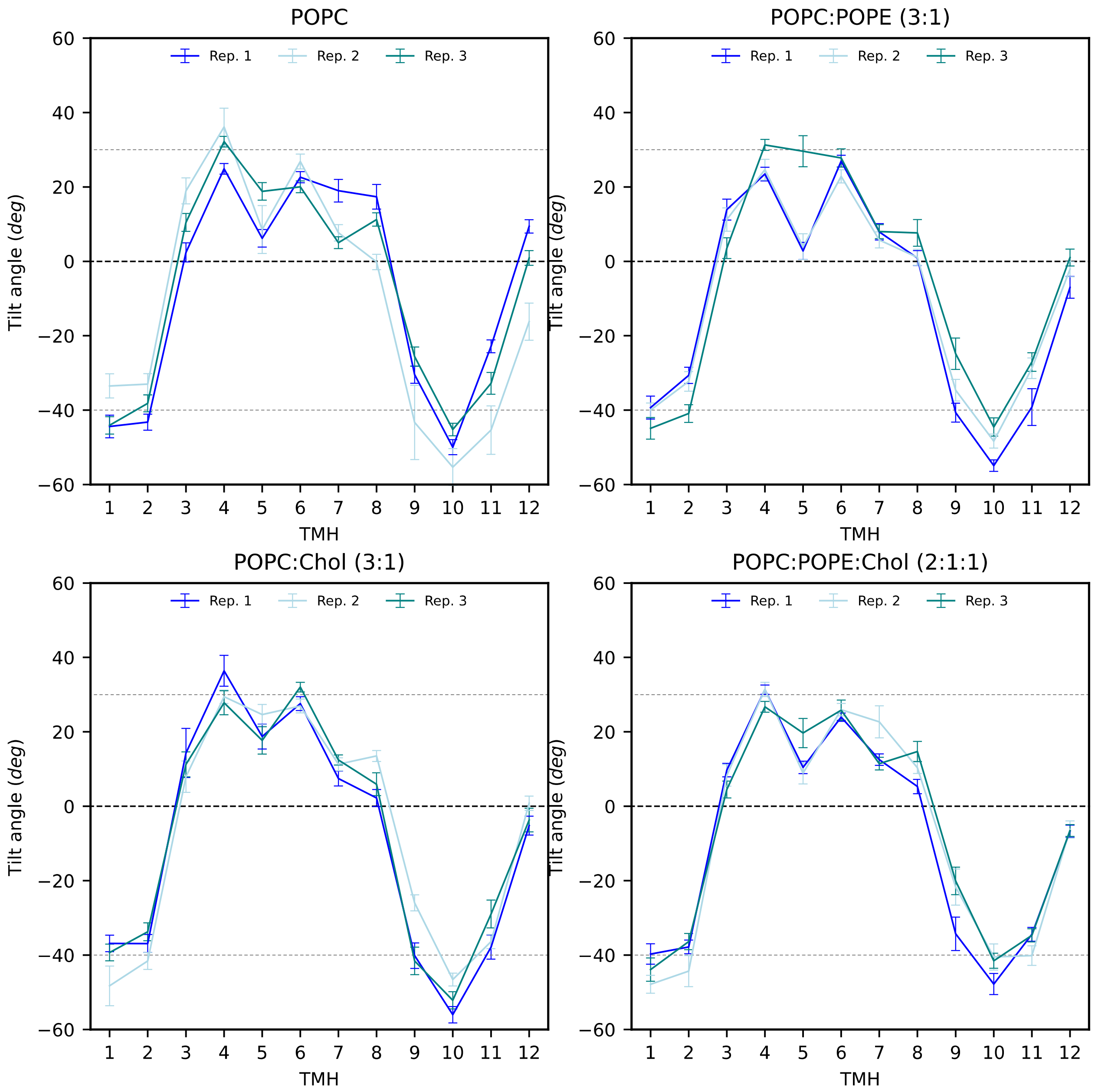

**Supplementary Figure S17.** The tilt angle profile (averaged over MD simulations) of *h*OAT1 (OF) in different lipid bilayers.

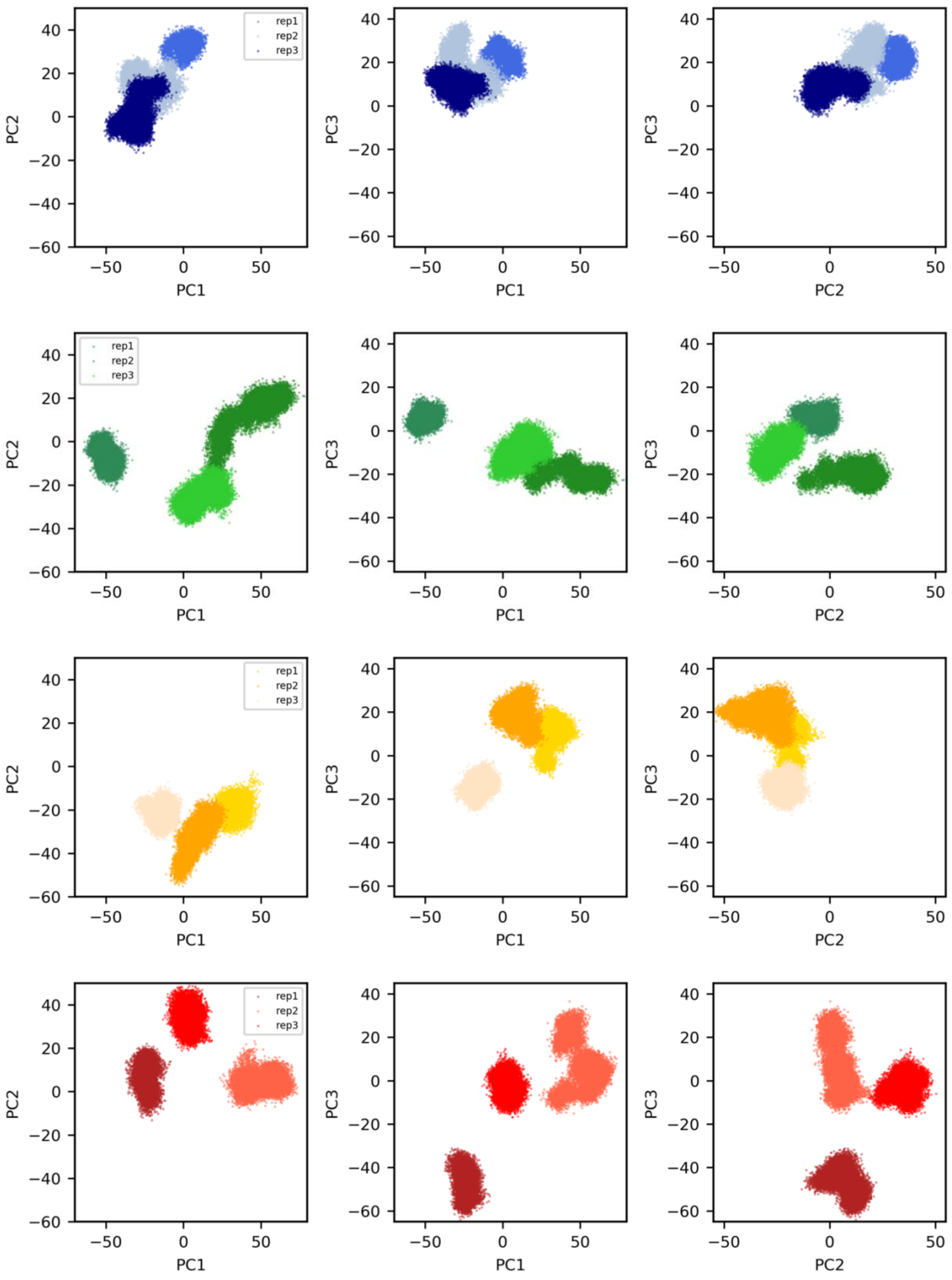

rep1
rep2
rep3
PC1
PC2
PC3

**Supplementary Figure S18.** PCA projected on all membrane conformational space of *h*OAT1 embedded in POPC:POPE:Chol (2:1:1) (blues), POPC:Chol (3:1) (greens), POPC:POPE (3:1) (yellows) and POPC (reds)

(A)

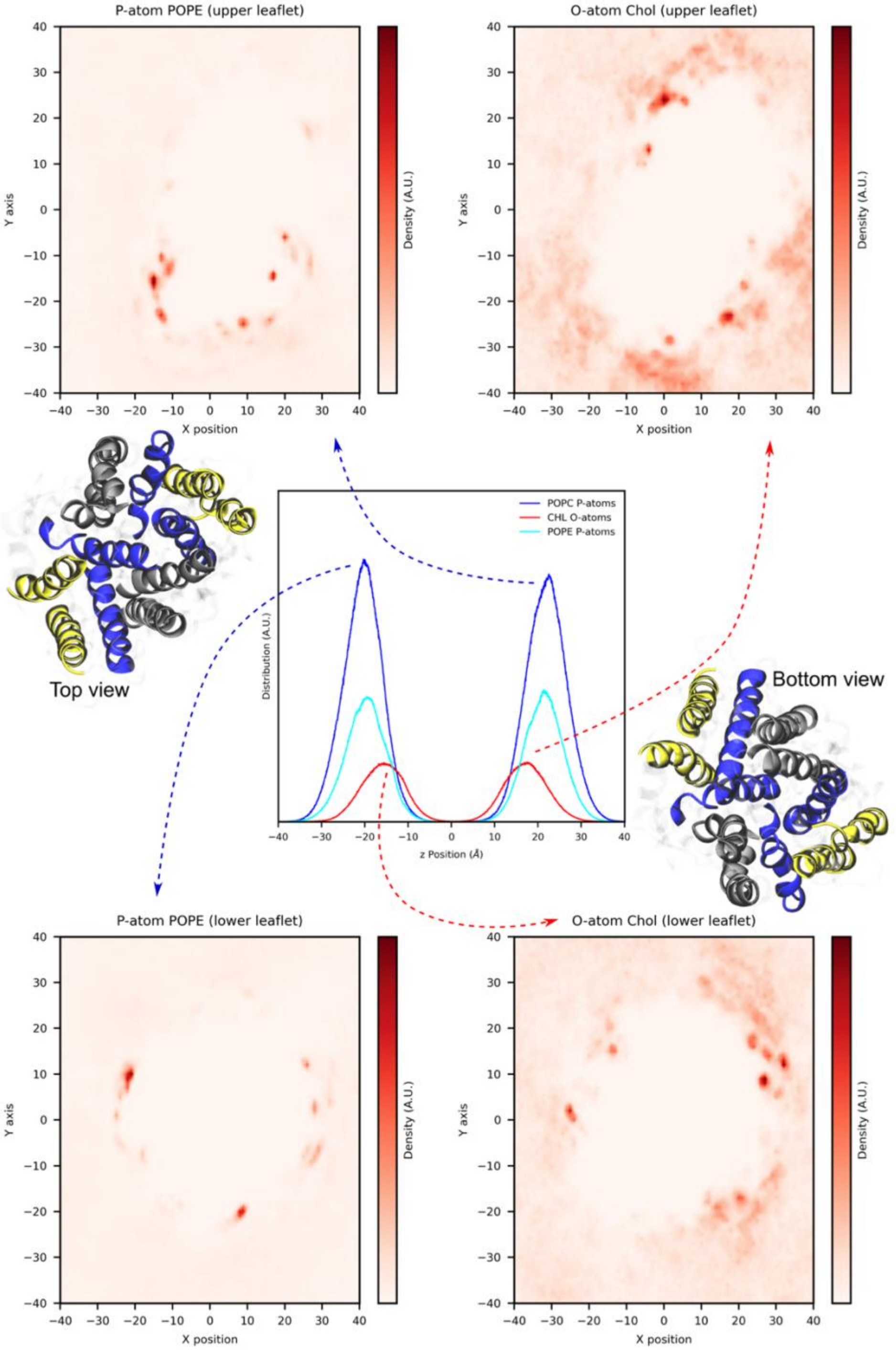

P-atom POPE (upper leaflet)
O-atom Chol (upper leaflet)
Y axis
X position
Density (A.U.)
POPC P-atoms
CHL O-atoms
POPE P-atoms
Distribution (A.U.)
z Position (Å)
Top view
Bottom view
P-atom POPE (lower leaflet)
O-atom Chol (lower leaflet)

(B)

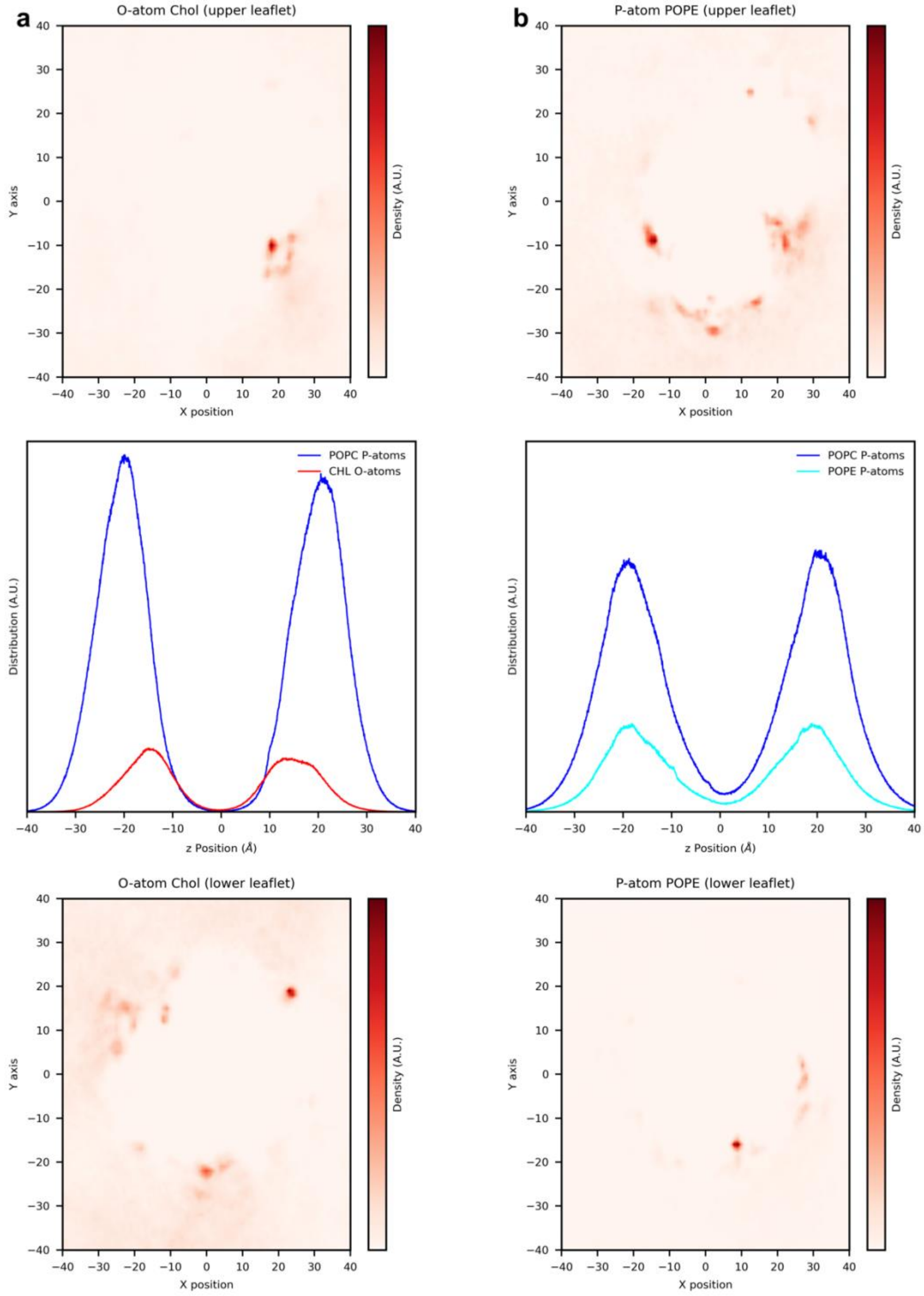

**Supplementary Figure S19.** Lipid density maps of *h*OAT1 inserted into (A) POPC:POPE:Chol (2:1:1). Upper and bottom leaflet of POPE (on the left) and cholesterol (on the right) are represented as density of P and O atom, respectively. (B) Lipid density maps of *h*OAT1 inserted into POPC:Chol (3:1) and POPC:POPE (3:1), shown with subpanel a and b, respectively.
